# An integrated mosquito small RNA genomics resource reveals dynamic evolution and host responses to viruses and transposons

**DOI:** 10.1101/2020.04.25.061598

**Authors:** Qicheng Ma, Satyam P. Srivastav, Stephanie Gamez, Fabiana Feitosa-Suntheimer, Edward I. Patterson, Rebecca M. Johnson, Erik R. Matson, Alexander S. Gold, Douglas E. Brackney, John H. Connor, Tonya M. Colpitts, Grant L. Hughes, Jason L. Rasgon, Tony Nolan, Omar S. Akbari, Nelson C. Lau

## Abstract

Although mosquitoes are major transmission vectors for pathogenic arboviruses, viral infection has little impact on mosquito health. This immunity is due in part to mosquito RNA interference (RNAi) pathways that generate antiviral small interfering RNAs (siRNAs) and Piwi-interacting RNAs (piRNAs). RNAi also maintains genome integrity by potently repressing mosquito transposon activity in the germline and soma. However, viral and transposon small RNA regulatory pathways have not been systematically examined together in mosquitoes. Therefore, we developed an integrated Mosquito Small RNA Genomics (MSRG) resource that analyzes the transposon and virus small RNA profiles in mosquito cell cultures and somatic and gonadal tissues across four medically important mosquito species. Our resource captures both somatic and gonadal small RNA expression profiles within mosquito cell cultures, and we report the evolutionary dynamics of a novel Mosquito-Conserved piRNA Cluster Locus (MCpiRCL) composed of satellite DNA repeats. In the larger culicine mosquito genomes we detected highly regular periodicity in piRNA biogenesis patterns coinciding with the expansion of Piwi pathway genes. Finally, our resource enables detection of crosstalk between piRNA and siRNA populations in mosquito cells during a response to virus infection. The MSRG resource will aid efforts to dissect and combat the capacity of mosquitoes to tolerate and spread arboviruses.

## INTRODUCTION

Mosquitoes are one of the most prevalent vectors of human pathogens, yet there is a wide variation in ability of the subclades to vector different pathogens. For example, all human malaria parasites are exclusively vectored by anopheline mosquitoes yet these same mosquitoes transmit few viruses other than O’Nyong nyong virus (ONNV) and Mayaro virus (Vanlandingham et al. 2006; Brustolin et al. 2018). On the contrary, culicine mosquitoes frequently transmit a range of human viral pathogens, such as dengue virus (DENV), Zika virus (ZIKV), Chikungunya virus (CHIKV) and yellow fever virus (YFV) in tropical climates where *AeAlbo* and *AeAeg* thrive; whereas eastern equine encephalitis virus (EEEV) and West Nile Virus (WNV) are spread mainly in *Culex* mosquitoes that inhabit more temperate climates (Conway et al. 2014; Olson and Blair 2015; Londono-Renteria and Colpitts 2016; Halbach et al. 2017; Lambrechts and Saleh 2019).

Since vector-pathogen interactions are complex and variable, no dominant theory yet explains why anopheline mosquitoes are less prolific than culicine mosquitoes in spreading arboviruses. Arbovirus infections in humans lead to devastating symptoms including fever, nausea, bleeding, extreme pain, brain damage and death. However, culicine mosquitoes showing highly active arbovirus replication are practically unaffected (Goic and Saleh 2012; Blair and Olson 2014; Olson and Blair 2015; Lambrechts and Saleh 2019) and therefore are highly competent transmitters of arboviruses to human hosts.

The three main classes of animal small regulatory RNAs are microRNAs (miRNAs) and endogenous small-interfering RNAs (endo-siRNAs), which range in size between 18-23nt long and are typically bound by Argonaute proteins; and Piwi-interacting RNAs (piRNAs) that are bound by Piwi proteins and which range in size between 24-32nt in length. Arguably, the most extensively characterized animal small regulatory RNAs are from *Drosophila melanogaster*, whose compact genome has decades of careful annotation and pioneering genetics tools. Many groups including ours have characterized *D. melanogaster* (*Dmel*) small RNAs comprising 258 miRNA genes (Kozomara et al. 2019), ∼20 large intergenic piRNA cluster loci also referred to as master control loci (Brennecke et al. 2007; Malone et al. 2009; Wen et al. 2014), >1000 genic piRNA cluster loci (Robine et al. 2009; Wen et al. 2014; Chirn et al. 2015), and >1000 endogenous loci generating either large fold-back transcripts or sense-antisense pairing transcripts that can give rise to endogenous siRNAs (Czech et al. 2008; Ghildiyal et al. 2008; Kawamura et al. 2008; Mirkovic-Hosle and Forstemann 2014; Wen et al. 2014; Wen et al. 2015). Finally, arbovirus-specific siRNAs and piRNAs are also detected in systemically infected *Dmel* cell cultures (Flynt et al. 2009; Wu et al. 2010; Vodovar et al. 2011; Goic et al. 2013; Wen et al. 2014; Palmer et al. 2018).

Culicidae mosquitoes are relatives of Drosophilid fruit flies as members of the Dipteran insect clade (**Figure 1A**, (Wiegmann et al. 2011)), yet ∼260 Million Years Ago (MYA) of evolutionary distance between Drosophilids and Culicidae imparts physiological and molecular differences in small RNA compositions. Within mosquito phylogeny, the anopheline subclade represented by *Anopheles gambiae* (*AnGam*) display a greater degree of chromosome synteny to Drosophilids than the culicine subclade of mosquitoes such as *Culex quinquefasciatus* (*CuQuin*), *Aedes aegypti* (*AeAeg*) and *Aedes albopictus* (*AeAlbo*) (Dudchenko et al. 2017). Indeed, *AnGam*’s genome (∼0.28GB) is as compact in size as *Dmel*’s genome (∼0.18GB), whereas culicine mosquito genomes are an order of magnitude greater in size due to massive accumulation of non-coding and repetitive elements (Fig. 1C) (Rai and Black 1999; Holt et al. 2002; Nene et al. 2007; Arensburger et al. 2010; Chen et al. 2015; Dudchenko et al. 2017; Matthews et al. 2018; Palatini et al. 2020).

**Figure 1.**
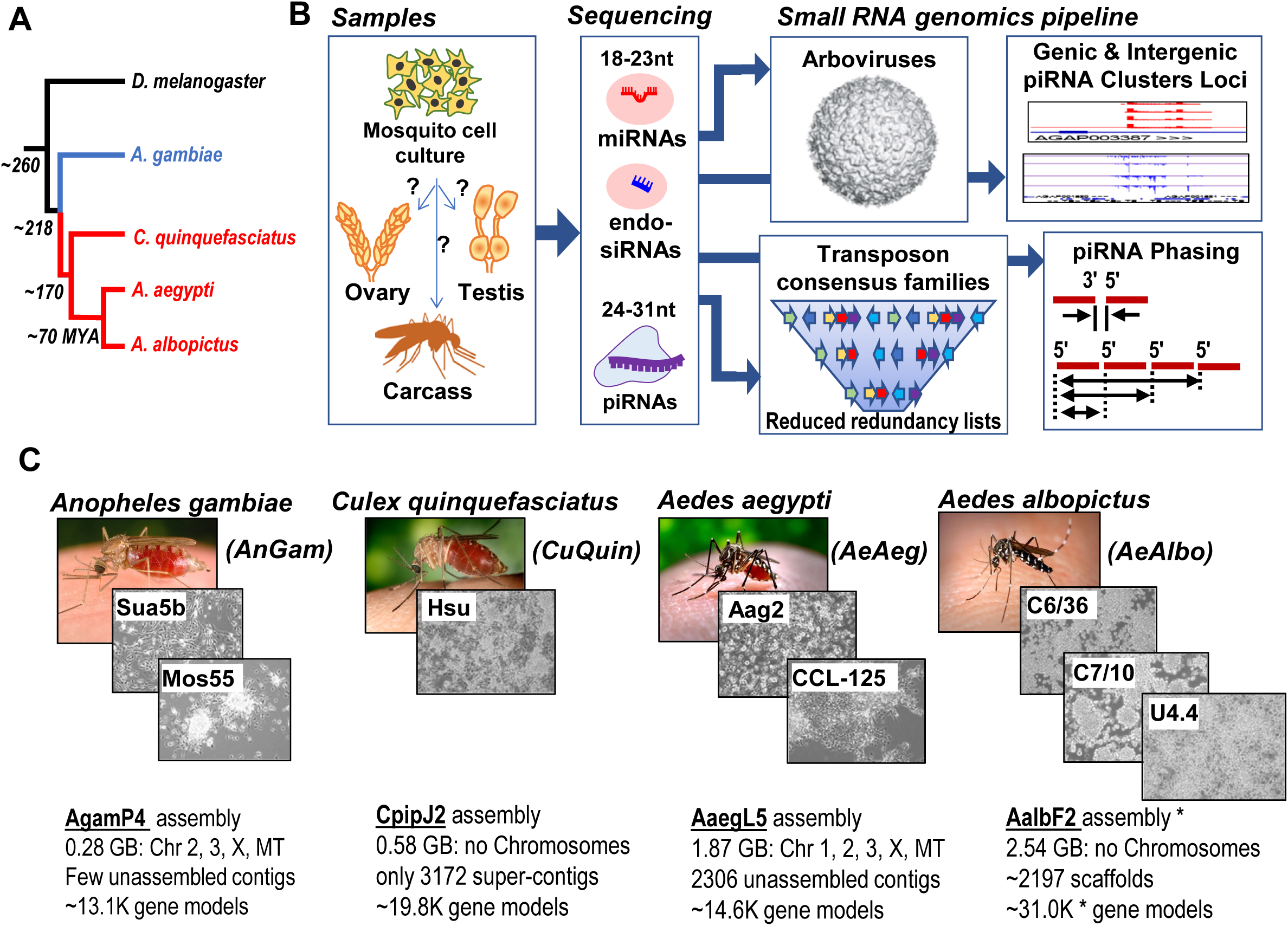
Overview of the mosquito small RNA genomics resource. (A) Phylogenetic tree of Dipteran insects in this study, with evolutionary distance measured by Million Years Ago (MYA). Blue and red color denote the anopheline and culicine lineages. (B) Organization of this resource that compares mosquito cell cultures to tissue types via determining the small RNA types and their genomic profiles. (C) Overview of the four mosquito species genomes and eight cell culture lines subjected to the small RNA genomics analysis pipeline. The specific genome assembly names are noted with genome configuration statistics below. The asterisk by the *AeAlbo* AalbF2 assembly indicates the early stage assembly annotation has a redundant list of gene models.

Since many viruses replicate their RNA genomes via a double-stranded RNA (dsRNA) intermediate, an antiviral mechanism conserved from plants to insects is the RNA interference (RNAi) pathway that uses Dicer and Argonaute enzymes to convert viral dsRNA into siRNAs that target viral mRNAs for gene silencing (Samuel et al. 2018; Guo et al. 2019). Recently, the piRNA pathway has also been implicated in assisting the siRNA pathway with an antiviral response in the culicine mosquitoes of *AeAeg*, *AeAlbo* and *CuQuin*, including in these species, cell culture lines (Goic and Saleh 2012; Blair and Olson 2014; Olson and Blair 2015; Halbach et al. 2017; Lambrechts and Saleh 2019).

A key knowledge gap in this field is to what degree viral siRNAs and piRNAs can comprise of or affect the endogenous small RNA transcriptome. Previous studies have examined mosquito small RNAs but have mainly focused on either virus derived small RNAs (Myles et al. 2008; Myles et al. 2009; Sanchez-Vargas et al. 2009; Brackney et al. 2010; Scott et al. 2010; Hess et al. 2011; Morazzani et al. 2012; Saldana et al. 2017; Varjak et al. 2017a; Varjak et al. 2017b; Ruckert et al. 2019); or conducted genomic analyses on earlier incomplete assemblies and preliminary annotations of individual mosquito species (Akbari et al. 2013; Whitfield et al. 2017; Tassetto et al. 2019).

In this study, we generated over 50 new small RNA libraries from mosquito cell culture lines, male and female gonads and respective carcasses from four medically important mosquito species (*AnGam, CuQuin, AeAeg, AeAlbo*) to add to the existing trove of publicly available small RNA libraries from previous mosquito lab studies. We then implemented our small RNA analysis pipeline tuned to the genome annotation features of each species and produced outputs amenable to cross-species comparisons. Our analysis provides the first holistic view of small RNA transcriptomes across mosquito phylogeny by revealing novel evolutionary and host dynamics in viral and somatic piRNA production and the strongest signal of periodicity in phased piRNA biogenesis patterns within culicine mosquitoes. Our efforts yield an improved comparative resource of Mosquito Small RNA Genomics (MSRG, http://class.bu.edu/~nclau/MSRG_db/MSRG_db_Home.html).

## RESULTS

### Framework for integrated small RNA analysis across four mosquito species

We previously built functional annotation pipelines for small RNA libraries generated from the gonads of Drosophilids, mammals and other vertebrates whose genome sequences and annotations were available in the UCSC genome browser (Chirn et al. 2015). Our original pipeline consisted of a series of shell, Perl and C scripts coupled with various short read mapping packages like Bowtie and Bowtie2 as well as BLAST and BLAT (Altschul et al. 1990; Kent 2002; Langmead et al. 2009; Langmead and Salzberg 2012). Together, the pipeline determines read length distributions, assigns reads to defined lists of miRNAs and structural RNAs such as transfer and ribosomal RNAs; then maps remaining reads to the genome with annotation overlays that allow for binning and counting of reads mapping to genes and predicted gene models, transposon consensus sequences, and intergenic regions. This pipeline was the first to enable comparative genomics of piRNA Cluster Loci (piRCL) between Drosophilids, mammals and other vertebrates; and showed that despite rapid evolution of these non-coding small RNA producing loci there exists a remarkable set of Eutherian-Conserved piRNA Cluster (ECpiC) loci with roles in mammalian reproduction (Chirn et al. 2015).

To improve our functional annotation pipeline for small RNAs in mosquito genomes (Fig. 1C), we added a curated list of arboviruses to the same plotting scripts as the transposon consensus sequences. We queried NCBI Genbank with search terms for mosquito arboviruses and viral genes from the literature (Nanfack Minkeu and Vernick 2018; Zakrzewski et al. 2018) and the Virus Pathogen Resource (VIPR)(Pickett et al. 2012), and arrived at a list of 225 mosquito arboviruses in May 2019 that exceeds the 107 Drosophilid viruses listed in (Palmer et al. 2018). Since Genbank includes multiple highly similar entries that are just slight sequence variants of a single virus class, we manually inspected literature and virus name entries to arrive at this curated list which we recognize will require future curation to keep current.

Finally, we took advantage of new genome assemblies of various culicine mosquito species coupled with additional genome annotation resources collected in the VectorBase database (Holt et al. 2002; Nene et al. 2007; Arensburger et al. 2010; Bartholomay et al. 2010; Giraldo-Calderon et al. 2015). *AeAeg* and *AeAlbo* genome assemblies have been enhanced with Hi-C information and longer reads sequencing to connect scaffolds into chromosomal assembles (Dudchenko et al. 2017; Matthews et al. 2018; Palatini et al. 2020).

### Reducing redundancy in mosquito transposon family consensus lists

Since *Dmel* makes abundant endo-siRNAs and piRNAs from transposon sequences, we needed to incorporate the mosquito transposon families’ set of consensus sequences into our small RNA annotation pipeline. *AnGam* and *Dmel* genomes have similar sizes and repetitive DNA amounts by biophysical determinations (Black and Rai 1988; Rai and Black 1999), yet the ∼3607 repeat families predicted for *AnGam* seemed excessive compared to ∼153 transposon families manually curated for *Dmel* (Kaminker et al. 2002). We suspected this redundancy in transposon family predictions was also rife in other mosquito genomes, with 3,730, 4,718, and 8,556 repeat families predicted in the *CuQuin*, *AeAeg*, and *AeAlbo* genomes, respectively (**Figure S1**). For perspective, mouse and human genomes are both larger than these mosquito genomes at ∼2.5Gb and ∼3.2Gb respectively, yet their curated lists in Repbase are ∼347 and ∼546 transposon families (Kapitonov and Jurka 2008).

Although it is possible there is more rapid evolution and therefore sequence diversity amongst transposon families in mosquitoes (Tu and Coates 2004; Goubert et al. 2017; Petersen et al. 2019), this feature cannot validate the order of magnitude greater number of transposon family entries which would lead to overestimation of transposon-mapping small RNAs. To reduce the redundancy in the *AnGam* repeats lists, we prioritized the curated Repbase entries and first used BLAT to merge other entries showing 90% sequence similarity to a Repbase entry (Fig. S1A, top). NCBI entries not merged to Repbase entries were then used as the second choice for centroids in the next iteration of BLAT, followed by EMBL entries and finally TE-fam entries. This process reduced the *AnGam* repeats to 555 transposon consensus sequences that helped us focus our small RNA analysis. This focused list is just ∼1/6^th^ of the original list, and we only reduced genomic coverage of transposons modestly <1%, yet we reduced the average double counting of small RNAs per *AnGam* library by ∼250% (Fig. S1A, bottom). We successfully applied similar but distinct redundancy reduction measures to transposon family consensus sequences lists for *CuQuin*, *AeAeg*, and *AeAlbo* as described further in the **Supplementary Text** and Fig. S1B-S1D.

### Anopheles gambiae (AnGam) soma have limited piRNAs whereas viral piRNAs are only detected in cell lines

We ran our small RNA genomics pipeline on published *AnGam* small RNA datasets as well as on new *AnGam* small RNA libraries we generated from Sua5b and Mos-55 cells originating from the Rasgon lab and an independent isolate of Mos-55 cells from the Colpitts lab, all later established in the Lau lab under culture conditions described in **Table S1**. We also generated new ovary and testis small RNA libraries matched to new libraries from the female and male carcasses devoid of gonads, and a selection of all the *AnGam* small RNA libraries we processed (**Table S2**) were scrutinized for the functional annotations of the small RNAs, the length distribution of the reads, and highlighting abundant small RNA mapping patterns to arboviruses persistently infecting the Sua5b and Mos55 cells (**Figure 2**).

**Figure 2.**
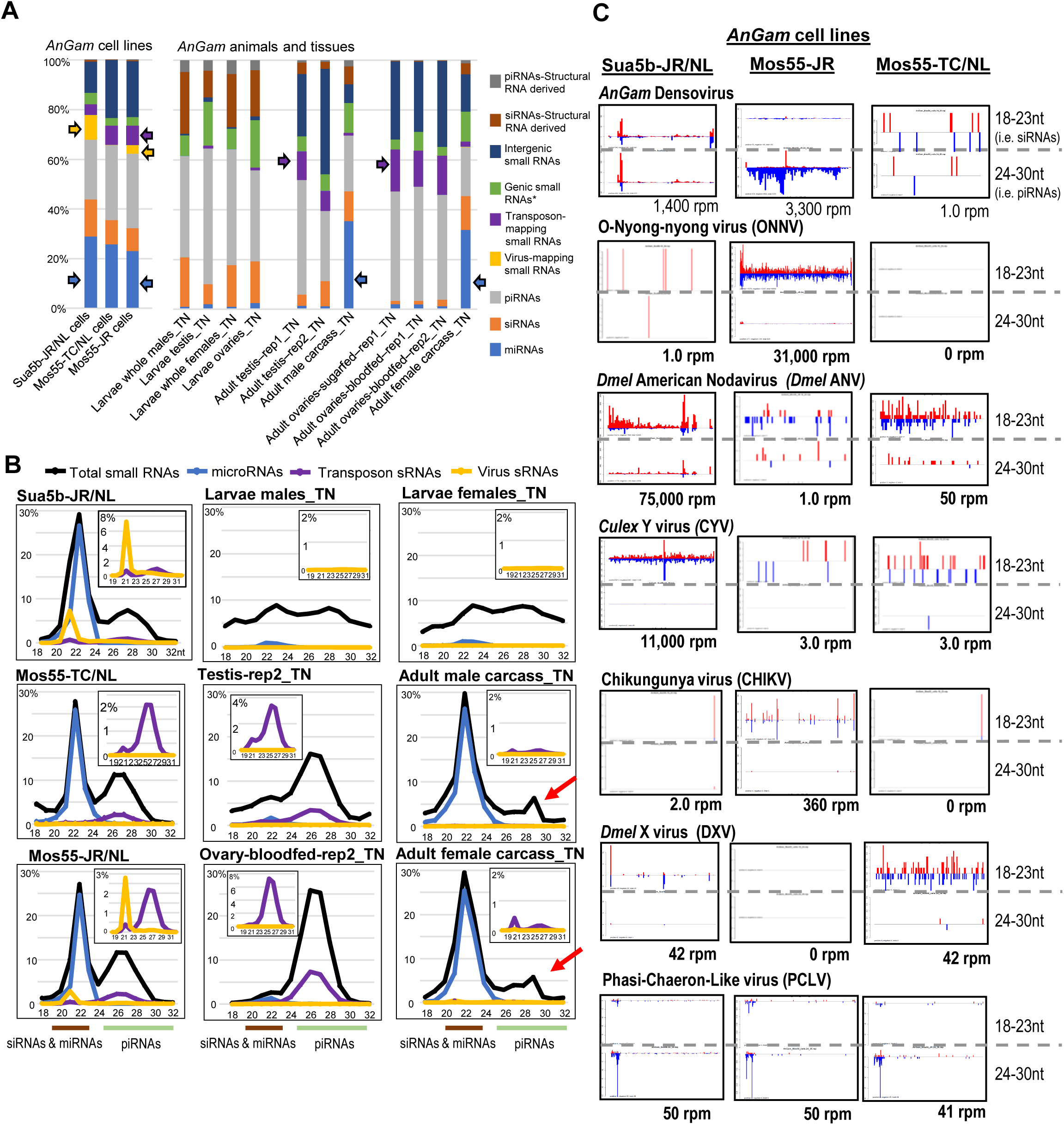
*Anopheles gambiae* (*AnGam*) small RNA profiles showing viral small RNAs only in *AnGam* cell lines. (A) Functional compositions of small RNAs in *AnGam* cells and tissues. Arrows point to notable portions of miRNAs, transposon-mapping small RNAs, and viral small RNAs. (B) Small RNA size distributions from *AnGam* cells and tissues, with colored lines marking the siRNAs and miRNAs ranging between 19-23nt, while piRNAs are between 24-30nt. The inset charts magnify the distribution of transposon and virus sRNAs under a different Y-axis range. (C) Profiles of viral small RNAs in *AnGam* cell culture lines. Reads per million (rpm) numbers are totals of the siRNA-length and piRNA-length small RNAs that come from the plus strand in red and minus strand in blue. The suffix to sample names is the initials of the laboratory investigator where the sample was originally obtained.

As expected, the bulk of *AnGam* gonad small RNAs were of piRNA lengths (24-32nt), whereas miRNAs were the major fraction in the carcasses (Fig. 2A, 2B). However, piRNAs and siRNAs were dominant over miRNAs even in the entire larvae. *AnGam* transposon-mapping small RNAs were also expressed highest in gonads, peaking at 8% of all the small RNAs, and are less than half of the proportional range of *Dmel* small RNAs mapping to transposon sequences (**Table S3**,∼18% in ovary). Both Sua5b and Mos55 cells expressed abundant miRNAs and piRNAs but the Mos55 cells displayed appreciably more transposon-mapping piRNAs that were similar to the profiles in gonads. To further compare these small RNA patterns, we performed hierarchical clustering by miRNAs and transposons counts, which mainly divided the small RNA libraries samples into the groups of gonads, carcass and larvae, and the cell lines (**Figure S2**).

To enable future studies of transposon regulation in mosquito cell culture lines that are to biochemistry and functional genomics approaches, we focused on highly abundant transposon small RNAs common to both cell lines and gonads, such as the small RNA profiles for *Tc1-1*, *Gypsy17*, *Gypsy60* and *Bel11* transposons (Fig. S2C). The transposon-mapping piRNAs dominated over siRNAs in both gonads and cell lines. However, there were also cell-line specific differences such as *Gypsy60* and *Bel11* that were only expressed highly in Mos-55 and Sua5b lines, respectively.

Similar to *Dmel*, we determined in *AnGam* numerous genic piRNA cluster loci and a smaller number of large intergenic loci. However the transposon insertion density was lower and fewer genic piRCLs in *AnGam* were biased at the 3’ UnTranslated Region (3’UTR)(Table S2), in line with previous studies (Biryukova and Ye 2015; Castellano et al. 2015). Even with our improved curation of Intergenic piRCLs in *AnGam* (Table S2) we could only determine a limited set of loci abundantly expressed in both gonads and cell lines which were analogous to the *Dmel Flamenco* piRCL that is abundantly expressed in OSS/OSC cells and the follicle cells of the *Dmel* ovary (Fig. S2D and S2E)(Lau et al. 2009; Saito et al. 2009). Only a few *AnGam* piRCL exhibited dense transposon insertions, and like *Flamenco*, the piRNAs are mainly biased on one genomic strand. However, we could not find an *AnGam* equivalent of the *Dmel* dual-strand *42AB* piRCL, which is extremely young evolutionarily and restricted to just the very closest relatives of *Dmel* (Chirn et al. 2015). Instead, we observed a few examples of divergent and convergent transcriptional units and one dsRNA locus generating piRNAs and siRNAs (Fig. S2E).

Although lab raised *AnGam* tissues lacked viral small RNAs, cultured cell lines carried persistent viral infections that generated proportionally more viral siRNAs than piRNAs (Fig. 2B). Sua5b cells produced striking amounts of viral siRNAs to *Culex* Y-virus (CYV) and *Dmel* American Nodavirus (*Dmel* ANV), whereas the Mos55-JR cells produce many viral siRNAs to ONNV, a virus related to CHIKV and known to be transmitted by *AnGam* (Vanlandingham et al. 2006; Rezza et al. 2017). An independent culture of Mos-55 cells from the Colpitts lab had very minor amounts of small RNAs mapping to *Dmel ANV, Dmel X-virus (*DXV*), and* Phasi-Chaeron-Like virus (PCLV)(Fig. 2C), seemingly suggesting it lacked persistent viral infection.

Both Sua5b-JR and Mos55-JR cells from the Rasgon lab produced sizable amounts of densovirus small RNAs, with much more densovirus piRNAs in the Mos55 line than Sua5b. Although a densovirus engineered to contain GFP was previously shown to be able to infect and replicate in Mos55 cell to express green fluorescence (Suzuki et al. 2015), more recently the Mos55-JR cells no longer recapitulated the engineered densovirus GFP expression (JL Rasgon, personal communication). The strong anti-sense bias of the densovirus piRNAs in Mos55-JR cells and densovirus siRNAs in Sua5b-JR/NL cells explains the current lack of GFP expression from the engineered densovirus. However, when we tried to verify that Mos55-TC/NL were negative for densovirus because they did not produce any viral small RNAs, we were surprised to detect as much densovirus viral RNA and the single-stranded DNA genome of densovirus in Mos55-TC/NL as Mos55-JR (**Figure S3**). Therefore, detecting viral small RNAs clearly reflects inherent viral infection status, but not vice versa.

### Few viral small RNAs in Culex quinquefasciatus (CuQuin) cells and mosquitoes

We possessed only one *CuQuin* cell line called Hsu cells (Hsu 1971), and it has been used for testing infections of arboviruses commonly transmitted by *CuQuin* such as WNV, which has been shown to generate predominantly viral siRNAs (Ruckert et al. 2019). We reanalyzed the Ebel lab dataset from (Ruckert et al. 2019) as well as our distinct isolate of Hsu cells which shared the production of piRNAs and siRNA from the Merida virus (**Figure 3**), but our isolate also had a trace of PCLV, *Culex* Densovirus, and CFAV. MicroRNAs dominated over piRNAs in Hsu cells, and only after WNV infection was there a significant accumulation of viral siRNAs. We also sequenced new testis, ovary and carcass small RNA pools (**Table S4**) and were struck by the significant amount of somatic piRNAs in the carcass as well as many 18-23nt small RNAs in the carcass that were not annotated *CuQuin* miRNAs. Although the Hsu cell miRNAs are well-accounted for, there could be potentially several novel *CuQuin* miRNA genes we could not characterize because this CpipJ2 genome assembly (Arensburger et al. 2010) is farthest behind in coverage and scaffolding size compared to the other three mosquitoes in this study.

**Figure 3.**
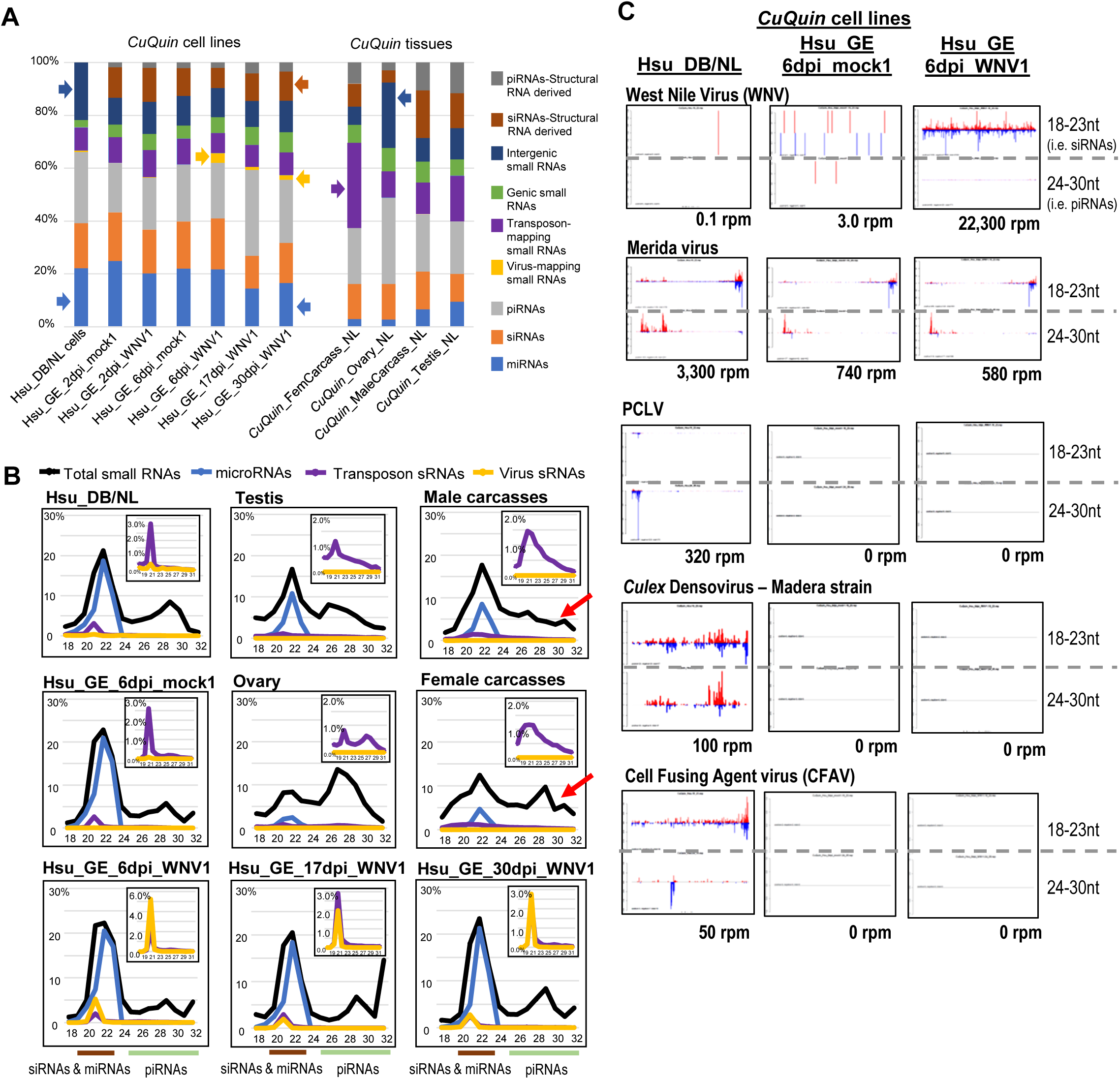
*Culex quinquefasciatus* (*CuQuin*) small RNA profiles are diminished in viral small RNAs. (A) Functional compositions of small RNAs in *CuQuin* cells and tissues. Colored arrows point to notable portions of miRNAs, transposon-mapping small RNAs, and viral small RNAs. (B) Small RNA size distributions from *CuQuin* cells and tissues, with colored lines marking the siRNAs and miRNAs ranging between 19-23nt, while piRNAs are between 24-30nt. The inset charts magnify the distribution of transposon and virus sRNAs under a different Y-axis range, and the red arrow points to low levels of somatic piRNAs. (C) Profiles of virus small RNAs in *CuQuin* cell culture lines. Reads per million (rpm) numbers are totals of the siRNA-length and piRNA-length small RNAs that come from the plus strand in red and minus strand in blue.

Amongst 1,700 *CuQuin* transposon families, the transposon small RNA mapping patterns are quite distinct between ovary and the female carcass (**Figure S4B**), whereas there was more common expression of transposon small RNAs between testis and male carcass. Hsu cells also expressed their own cell-line-specific set of transposon small RNAs, yet share more notable transposon small RNAs in ovary compared to the testis (Fig. S4B, S4C), since Hsu cells were derived from *CuQuin* ovaries (Hsu 1971). Examples of both retrotransposons and DNA transposons exhibited various distinct small RNA mapping patterns on both plus and minus strands (Fig. S4C). In contrast to *AnGam* piRNAs mostly mapping to a single genomic strand in a piRCL, *CuQuin* piRCL exhibit much more overlapping piRNAs on both genomic strands, including across regions containing genes (Fig. S4D) as well as intergenic regions (Fig. S4E). Interestingly, some of these intergenic piRCL contain multiple tandem large repeats, one identified by RepeatMasker as containing multiple iterations of a transposon, and a third locus missed by RepeatMasker but evidently displays tandem repeats (Fig. S4E). These instances of satellite DNA repeats serving as piRCL will be discussed further below.

### Persistent arbovirus infections in Aedes aegypti (AeAeg) cells and animals

The yellow-fever mosquito, *A. aegypti*, is a prolific vector for YFV, DENV, ZIKV, Mayaro virus, and other viral pathogens that exert a great toll on human health in tropical regions (Conway et al. 2014; Olson and Blair 2015; Londono-Renteria and Colpitts 2016; Halbach et al. 2017; Lambrechts and Saleh 2019). With our pipeline we could now annotate many published *AeAeg* small RNA datasets from many labs for comparison (Campbell et al. 2008; Myles et al. 2008; Hess et al. 2011; Adelman et al. 2012; Vodovar et al. 2012; Akbari et al. 2013; Liu et al. 2015; Miesen et al. 2016; Samuel et al. 2016; Dietrich et al. 2017; Saldana et al. 2017; Varjak et al. 2017a; Varjak et al. 2017b; Lewis et al. 2018). To examine biological variability, we also established our own cultures of two *AeAeg* cell lines, CCL-125 and Aag2 cells grown under conditions detailed in Table S1, as well as obtained ovary, testis, female and male carcasses from additional *AeAeg* isolates, each sample coded with the initials of the lab principal investigator (i.e. KM-Kevin Myles, BH-Bruce Hay, ZT-Zhijian Tu, FJ-Francis Jiggins, GH-Grant Hughes, TC-Tonya Colpitts, etc.). Small RNA libraries were sequenced and functionally annotated in **Figure 4** and **Table S5,** and in *AeAeg* cell cultures miRNAs were a much lower proportion of the total small RNA composition compared to *AnGam* and *CuQuin* cell cultures and consistent with *AeAeg* larvae and male and female carcasses.

**Figure 4.**
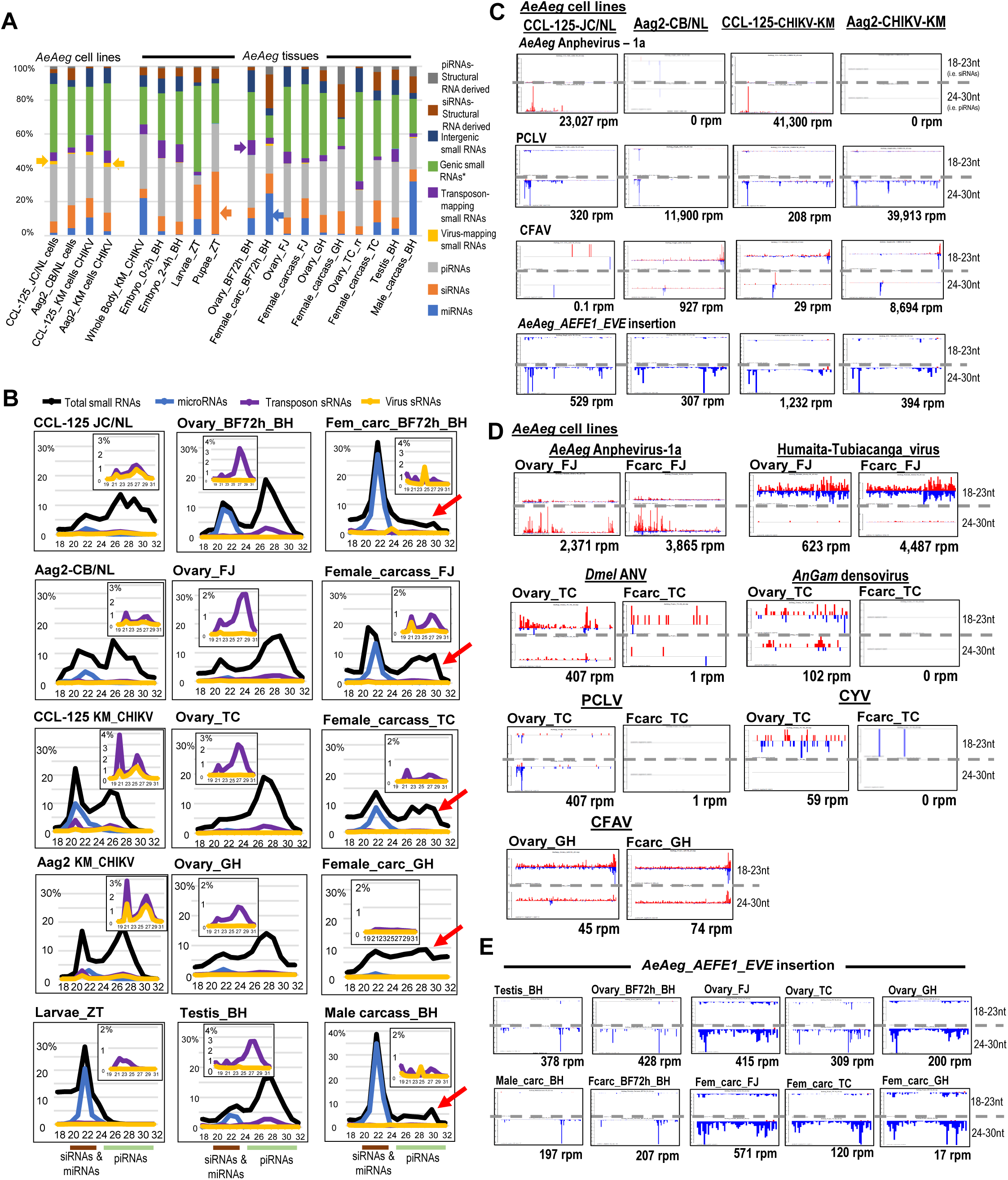
*Aedes aegypti* (*AeAeg*) small RNA profiles shows cells and mosquito isolates have distinct sets of persistent viral infections. (A) Proportions of small RNA functional annotations. Arrows point to notable portions of miRNAs, transposon-mapping small RNAs, and viral small RNAs. (B) Length distributions of small RNAs as a percentage of the library. Inset graphs zoom in on the proportions of transposon and virus sRNAs. Red arrows highlight data points of significant somatic piRNAs. (C) Mapping patterns and counts of notable viruses generating small RNAs in mosquito cells. (D) Persistent viral piRNAs in certain *AeAeg* strains that also exhibit greater amounts of somatic piRNAs, whereas the *AeAeg* strain from the Akbari et al (BH) study lacks inherent viral piRNAs and low amounts of somatic piRNAs. (E) Profiles and counts of mainly antisense piRNAs from the Endogenous Virus Element (EVE) AEFE1 / AY347953.2 that has homology to the NS5 gene of flaviviruses like CFAV.

Making up the bulk of the *AeAeg* cell lines’ small RNAs are piRNAs within intergenic regions as well as a significant proportion overlapping genic regions in large swaths across many genes (i.e. **Figure S5D**, leftmost snapshot, and Table S5). In contrast to *Drosophila* and mammalian genic piRCL (Chirn et al. 2015), only a small handful of *AeAeg* genic piRCL had configurations suggesting the main precursor transcript is the gene mRNA’s 3’UTR (Fig.5D middle and right snapshots). Similarly, few 3’UTR genic piRCL were observed in *AnGam* and *CuQuin*. Although there are many scattered transposons throughout the *AeAeg* genome with transposon-mapping piRNAs, we curated *AeAeg* intergenic piRCL to the largest contiguous loci (Table S5) with examples in Fig. S5E highlighting a piRCL with a single-genomic strand bias, expression from both genomic strands suggesting convergent transcription, and another example of a piRCL with satellite tandem repeats. Overall, the density of transposon in *AeAeg* piRCL were low as reflected also by the minor fraction overall of transposon-mapping small RNAs (Fig. 4A, 4B). Although gonads expressed the highest level of transposons small RNAs, some of which are also highly expressed in *AeAeg* cell lines (Fig. S5B), these transposon-mapping small RNAs are less likely to originate from piRCL but perhaps from other single transposons loci, as suggested in *Dmel* (Shpiz et al. 2014; Senti et al. 2015).

**Figure 5.**
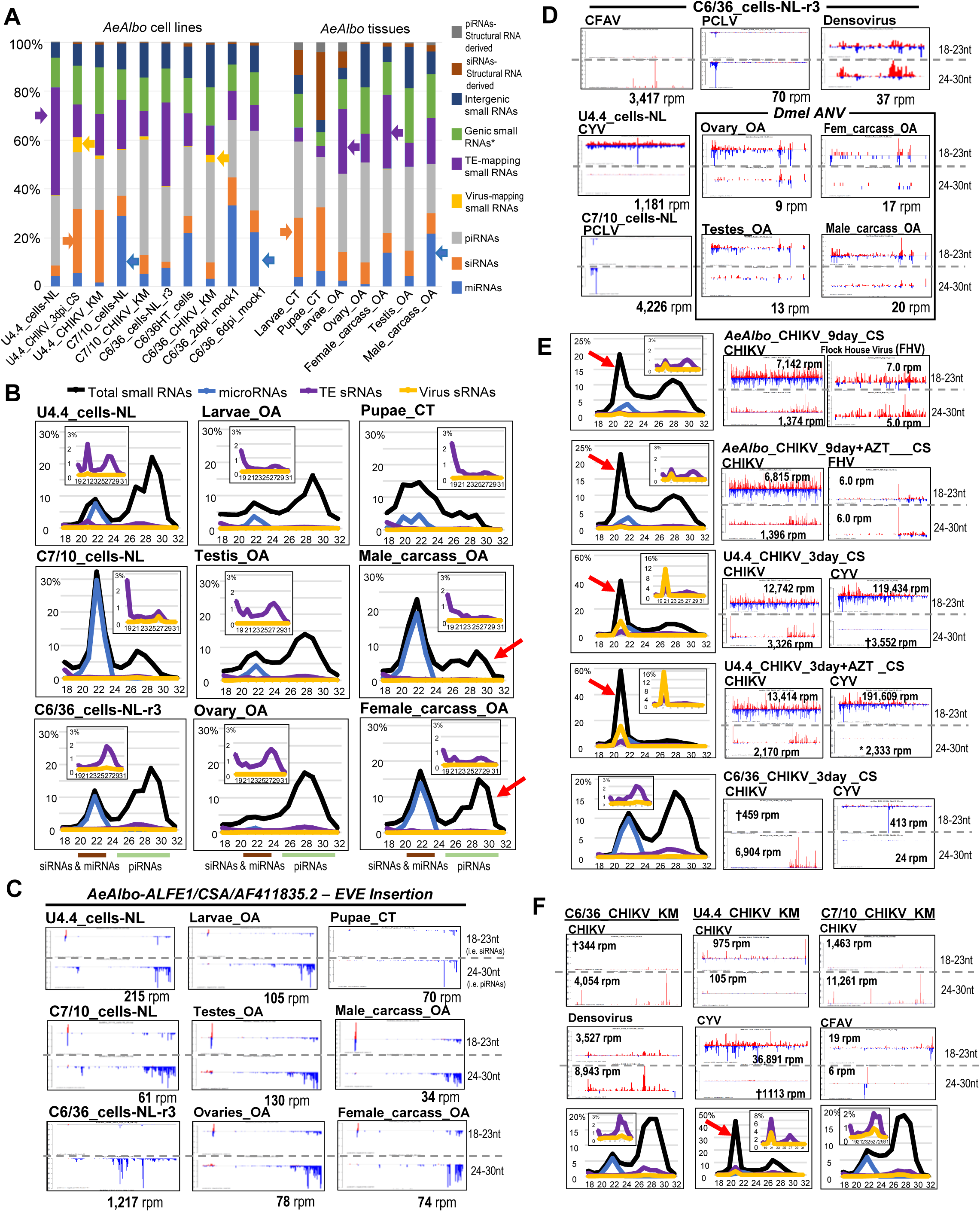
*Aedes albopictus* (*AeAlbo*) cells and animals are prolific in persistent viral small RNAs and somatic piRNAs. (A) Functional compositions of small RNAs in *AeAlbo* cells and tissues. Arrows point to notable portions of miRNAs, TE-mapping small RNAs, and viral small RNAs. (B) Small RNA size distributions from *AeAlbo* cells and tissues, with brackets highlighting siRNAs and miRNAs between 19-23nt, while piRNAs are between 24-30nt. The inset charts magnify the distribution of TE and virus sRNAs under a different Y-axis range. (C) Gallery of small RNAs from an Enodgenous Viral Element in the *AeAlbo* genome, with homolog to capsid and NS5 proteins of flaviruses like CFAV. Reads per million (rpm) numbers are totals of the siRNA-length and piRNA-length small RNAs that come from the plus strand in red and minus strand in blue. (D) Inherent viral small RNAs within the libraries sequenced in this study. (E) Small RNA size distributions and profiles and counts of *AeAlbo* mosquitoes and cells infected with CHIKV from the Saleh lab; (F) and for comparison *AeAlbo* cells infected with CHIKV from the Myles lab. A top persisting arbovirus generating the most viral small RNAs is shown next to CHIKV, whereas red arrows point to the massive proportion of siRNAs in *AeAlbo* mosquitoes and U4.4 cells from the Saleh and Myles labs. † marks small RNA reads that appear to be squashed by the Y-axis of the other plot in that diagram.

Despite the wide competency of *AeAeg* cells and mosquitoes to support arbovirus replication, viral piRNAs made up a small proportion of the total small RNA populations, even with an ectopic infection of CHIKV, DENV or ZIKV (<∼6% Table S4-tab Metatable, Fig. 4A, 4B). *AeAeg* mosquitoes and cell cultures are known to remain phenotypically unaffected by arbovirus infection presumably because of antiviral RNAi pathways that generate viral siRNAs and piRNAs, as described previously by many labs (Aliyari and Ding 2009; Karlikow et al. 2014; Blair and Olson 2015; Samuel et al. 2018). Our annotation pipeline also confirmed earlier findings of viral piRNAs and persistent viral infections of PCLV and CFAV in CCL-125 and Aag2 cells (Fig. 4C and **Figure S6A**), but we also discovered new additional viral piRNAs in the *AeAeg* cell cultures and certain *AeAeg* isolates (Fig 4C, 4D and Fig. S6A). We discovered that *AeAeg* Anphevirus strain-1a piRNAs are robustly generated in CCL-125 cells as well as in the ovary and soma of just one wild *AeAeg* isolate from Miami, USA (Lewis et al. 2018), samples Ovary_FJ and Fcarc_FJ) but absent in other *AeAeg* isolates and Aag2 cells.

Whereas Anphevirus piRNAs strongly dominated over siRNAs, viral siRNAs were the dominant response to a Humaita-Tubiacanga virus in the Miami/Jiggins-lab *AeAeg* isolate (Lewis et al. 2018), *Dmel* ANV siRNAs mainly in ovary of an *AeAeg* isolate from the Colpitts-lab, and CFAV siRNAs and piRNAs in both ovary and soma of an *AeAeg* isolate from the Hughes-lab (Fig. 4D) (Saldana et al. 2017).

### Variability in the levels of mosquito somatic piRNAs

A main function of the Piwi pathway and piRNAs in *Drosophila* and mammals is to silence transposons in gonads to ensure fertility, with much less evidence for somatic functions because of at least a magnitude less expression of somatic piRNAs (Lewis et al. 2018; Genzor et al. 2019). However, *Drosophila* is actually an outlier in the insect class with respect to a lack of expression of Piwi pathway genes and somatic piRNAs (Lewis et al. 2018), whereas mosquitoes are capable of expressing significant amounts of somatic piRNAs. Some of these endogenous piRNAs were recently proposed to have an antiviral role by being complementary to flavivirus sequences but were primarily generated from genomic insertions called Endogenous Viral Elements (EVEs, (Lequime and Lambrechts 2017; Suzuki et al. 2017; Whitfield et al. 2017; Houe et al. 2019; Tassetto et al. 2019; Blair et al. 2020)). The most active EVE in our dataset, the AEFE1/AY347953 EVE has homology to the NS5 gene of flaviruses like Kamiti River virus and CFAV (Crochu et al. 2004), predominantly generated piRNAs with a smaller amount of siRNAs in the gonads and soma (Fig. 4E) as well as in cell lines (Fig. 4C.). In contrast, the consistently antisense piRNAs to PCLV, largely from the S-fragment of the PCLV genome (left most region of the plot) suggests this is also an EVE signature (Whitfield et al. 2017; Tassetto et al. 2019). Although piRNAs from the CFAV-like EVE should theoretically target CFAV (Suzuki et al. 2017; Whitfield et al. 2017), there is still sufficient persisting replication of CFAV RNAs in the Hughes lab *AeAeg* isolate (Fig. 4D) and *AeAeg* cell cultures (Fig. 4C) to produce significant other CFAV small RNAs. We were unable to further cross-reference other analyses of *AeAeg* EVEs (Whitfield et al. 2017; Tassetto et al. 2019) because this was only performed on an incomplete genome assembly from their isolate of the Aag2 cell line.

However, we sought to examine the significant variability in some mosquito carcasses showing robust amounts of somatic piRNAs (i.e. Fig. 4B, *AeAeg* Female carcass_FJ, *_TC, *_ GH; and Fig. 3B, *CuQuin* Female carcass_NL) in contrast to others with very subdued somatic piRNAs (i.e. Fig. 4B, *AeAeg* Female carcass_BH, Male carcass_BH; Fig. 2B, *AnGam* Male carcass_TN, Female carcass_TN; and Fig. 3B, *CuQuin* Male carcass_NL). First, we conducted considerable inspection of our experimental and bioinformatics methods to rule out potential sources of unintended detection bias like residual gonads contaminating carcass and found none since no germline-specific transcripts like *vasa* were detected (data not shown). Then we hypothesized that the reason those three *AeAeg* isolates were expressing abundant somatic piRNAs is because they were also persistently infected by an arbovirus as reflected by viral small RNAs. This hypothesis is also supported by the absence in our analysis of viral small RNAs in the *AeAeg* isolate from the Hay lab (Akbari et al. 2013) and the *AnGam* isolate from the Nolan lab (this study and (Castellano et al. 2015)).

### Potential crosstalk between piRNAs targeting distinct flaviviruses

To further explore our hypothesis that *AeAeg* piRNAs could be responding to arboviral infections, we reanalyzed a well-matched small RNA dataset of female *AeAeg* mosquitoes fed blood that lacked or contained ZIKV, and reconfirmed that both ZIKV siRNAs and piRNAs only accumulated to detectable levels at 7 and 14 days post-infection (Saldana et al. 2017). Whereas bulk overall small RNA levels largely remain the same whether the mosquitoes are harboring ZIKV or not (Fig. S6B), our analysis revealed an interesting production of new piRNAs from a specific region of CFAV only stimulated after ZIKV replication (blue arrows in Fig. S6B). This region did not have specific homology to ZIKV piRNAs but generated both plus and minus strand piRNAs indicative of the “ping-pong” mode of piRNA interactions. Despite clear signals of ZIKV and CFAV small RNAs, these viral small RNAs were only a tiny fraction of the total small RNAs samples in these libraries (Fig. S6B, right most length distribution plots).

Therefore, to see if we could augment the proportion of exogenous arboviral small RNAs in mosquito cells during a response to infection, we conducted our own infection of Aag2 cells with DENV and ZIKV, which we performed in two independent infection experiments on different batches of Aag2 cells each of the four DENV serotypes, including two strains of the DENV2 serotype (NGC-a high passage and K0048-low passage) (Troupin et al. 2016); as well as the Old World (OW) and Puerto Rico (PR) isolates of ZIKV (Araujo et al. 2020) (Fig. S6C). Additional details on how we assessed viral infections of Aag2 cells are discussed in the Supplementary Text.

Although both batches of Mock Control Aag2 cells had expected bimodal distributions of 18-23nt siRNAs and miRNAs versus 24-32nt piRNAs, we observed instances were these distributions were greatly affected by viral infection. In both replicates, DENV2K0048 distorted these two distributions, in one case greatly enhancing endogenous siRNAs while depressing piRNAs, and in the other case a vice versa response (Fig. S6D and S6E, red arrows). Also, in both replicates, the ZIKV_OW infections enhanced endogenous siRNAs while depressing piRNAs, while this was vice versa in one ZIKV_PR infection. Although DENV2NGC, the high passage strain, repeatedly lacked impact on small RNA populations, there was marked variability in one of the experiments but not in the other for when DENV1, DENV3 and DENV4 infections greatly affected the bimodal distribution of piRNAs versus siRNAs and miRNAs. Besides confirming viral infection of all Aag2 cell samples by qRT-PCR (Fig. S6C), we could also readily detect the siRNAs and piRNAs to the specific DENV serotype (Fig. S6F). Whereas DENV and ZIKV siRNAs were generated from both plus and minus strands indicative of a dsRNA precursor, the viral piRNAs were biased from the plus strand and predominantly arising from a small set of very abundant reads (Fig. S6F bottom plots), recapitulating the same confounding patterns observed by others (Goic et al. 2016; Miesen et al. 2016; Whitfield et al. 2017; Merkling et al. 2020). This pattern of viral piRNA accumulation defies the generalized biogenesis patterns of phased piRNAs (Han et al. 2015; Mohn et al. 2015; Pandey et al. 2017; Gainetdinov et al. 2018; Izumi et al. 2020).

Future studies will be needed to dissect this experimental variability in Aag2 cells’ small RNA populations during arbovirus infection, but we revisited the Aag2 CFAV small RNA patterns since these CFAV piRNAs were altered in the Galveston strain of *AeAeg* mosquitoes after ZIKV infection (Fig. S6B). Because Aag2 cells are already persistently infected by three other arboviruses, the two batches of Mock Control cells already displayed an enhanced population of minus-strand piRNAs similar to the region of CFAV piRNAs amplified in the ZIKV-infected mosquitoes (Fig. S6G). DENV and ZIKV infections did not clearly affect these piRNAs, but it appears that these CFAV piRNAs being influenced by other viral infections corresponds to the NS2A gene (Fig. S6H). What specifies the NS2A gene as a piRNA precursor and CFAV 3’UTR as a stronger initiator of siRNA biogenesis remains unclear (Fig. S6H), although other flavivirus 3’UTRs have been described to have an antiviral role (Moon et al. 2015).

### Aedes albopictus (AeAlbo) cells and mosquitoes are widespread in producing viral and somatic piRNAs

*AeAlbo* rivals *AeAeg* as a prolific arbovirus vector because of its increased geographical range, more robust diapause stages, and greater tolerance of different climates (Gamez et al. 2020; Palatini et al. 2020). Three distinct *AeAlbo* cell lines have been frequently used to investigate arbovirus infections: U4.4, C7/10, and C6/36 cells. As determined previously, the C6/36 cell line is exceptional for having mutations that inactivate its *Dicer2* gene so that it lacks robust RNAi activity, yet it robustly produces piRNAs (Scott et al. 2010; Weger-Lucarelli et al. 2018). Sequencing our own cultures of U4.4, C7/10 and C6/36 cells confirmed abundant piRNAs and rather low levels of siRNAs because miRNAs that made up the bulk of 18-23nt small RNAs (**Figure 5A, 5B**). We also sequenced small RNAs from ovary, testis and female and male carcass from an Akbari lab *AeAlbo* isolate from Los Angeles and discovered just as much somatic expression of piRNAs as in the gonads (Fig. 5B, **Table S6**). Since the other mosquito carcasses that had low amounts of somatic piRNAs also lacked a persistent virus that we could detect from viral small RNAs, we hypothesized *AeAlbo* animals and cells would contain persistent viruses.

We first detected in both *AeAlbo* tissues and cell culture lines abundant piRNAs mapping antisense to an *AeAlbo* EVE insertion with various names like ALFE1, CSA, and AF411835.2 (Fig. 5C, (Crochu et al. 2004; Suzuki et al. 2017)), which like the similar EVE in *AeAeg*, AEFE1 (Fig. 4E), has homology to CFAV NS5 protein but does not absolve *AeAlbo* cell cultures from expressing persistent CFAV small RNAs (Fig. 5D, 5F). We detected various other viruses generating persistent viral small RNAs, from PCLV to CYV to densovirus; and interestingly the densovirus piRNAs contributed to siRNA-length small RNAs from two independent C6/36 cell isolates (Fig 5D, 5F) despite this cell line lacking *Dicer2* activity (Scott et al. 2010; Weger-Lucarelli et al. 2018). The CYV small RNAs were uniquely reserved to the U4.4 cell line and biased to generating siRNAs versus piRNAs, both in our lab’s line but also in the Myles and Saleh labs line (Morazzani et al. 2012; Goic et al. 2016) where the endogenous siRNA levels are exorbitant at 40%-60% of the total small RNAs (Fig. 5E, 5F). Although siRNA production in the Akbari lab *AeAlbo* mosquito was much lower compared to gonad and somatic piRNAs (Fig. 5B), the siRNA levels in the Saleh lab *AeAlbo* strain was also surprisingly high at >25% of the total small RNAs (Fig. 5E). However, it is unclear if this is because these *AeAlbo* mosquitoes are responding to acute CHIKV infection that predominantly generates siRNAs from both plus and minus strands, whereas CHIKV piRNAs are biased for production from the virus plus strand (Fig. 5E).

Multiple *AeAlbo* isolates displayed persistent viral infections such as *Dmel* ANV in the Akbari lab isolate and Flock House Virus (FHV) in the Saleh lab isolate (Fig 5D, 5E), perhaps because these labs utilized *Drosophila* in their studies and *Dmel* is a prominent reservoir for these two nodaviruses (Goic et al. 2013; Kandul et al. 2019). In addition, we also detected viral small RNA signatures like densovirus and ONNV from these reports (Morazzani et al. 2012; Wang et al. 2018), respectively (data not shown). However, with the exception of the U4.4 cells, viral small RNAs in *AeAlbo* were also a very minor fraction of the total small RNAs produced in these mosquitoes.

*AeAlbo* transposon-mapping small RNAs were also a minor but larger proportion comprised more by piRNAs than siRNAs and were also most highly expressed in the ovaries and testes (Fig. 5B, **Figure S7**). Although the *AeAlbo* cell cultures we examined expressed piRNAs, the hierarchical clustering of miRNA and transposon-mapping small RNA profiles still placed cell cultures together in their own group distinctly apart from either gonads or somatic tissues (Fig. S7A, S7B). Nevertheless, we could highlight several transposons commonly generating abundant piRNAs across *AeAlbo* cells and gonads, with a biased number of these examples as members of the LTR-class of transposons (Fig. S7C). This may be due to the new transposon consensus lists generated from RepeatModeler v2 on the AalbF2 assembly and >60% of the repeats it called were in the LTR-class.

In trying to define the transcriptional units of piRCL in this new AalbF2 genome assembly, we repeated manual curation of genic and intergenic piRCL which again deviated from known configurations in *Drosophila* and mammals (Chirn et al. 2015). A genic piRCL with very abundant piRNAs in the 3’UTR of the *AeAlbo* gene “XM_019699801” (Fig. S7D, leftmost window) only has a very sharp peak in the middle of the 3’UTR in contrast to the broad peaks of conventional genic 3’UTR piRCL, and interestingly is the ortholog of the *AeAeg* gene “AAEL01051” (Fig. S5D, rightmost window) that is also a genic 3’UTR piRCL. The six notable piRCL we highlighted exist on three super-scaffolds, with mostly single-stranded biases in the small RNA expression patterns or a gigantic 2.5Mb region holding 6 intergenic piRCL (Fig. S7E). Finally, we note two genic piRCL with divergent transcriptional directions reminiscent of mammalian intergenic piRCL, yet the patterns of these piRNAs display a pattern of satellite DNA repeats. (Fig. S7D, rightmost window).

### A major AeAlbo genic piRCL is dynamically evolving yet syntenically conserved through mosquito phylogeny

In the culicine mosquitoes of *CuQuin*, *AeAeg* and *AeAlbo*, we noted large intergenic piRCL with satellite DNA repeats generating very abundant amounts of piRNAs, but the lack of synteny made it challenging to compare these particular piRCL across the species. However, there was one additional piRCL with satellite DNA repeats that was linked to the *AeAlbo* genes “XR_003896043.1” and “XM_019702868.2” which have descriptions as a pro-resilin-like isoform or an isoform with similarity to the RNA-binding protein FUS (**Figure 6**). Expressed very highly in *AeAlbo* gonads, somatic tissues, and cell cultures, this genic piRCL generates an average >10,000 reads per million (rpm) from mainly two major piRNAs which have 33 and 27 alternating repeats spread out in a ∼5.6kb region (Fig. 6C). The *AeAeg* orthologous gene also contained a genic piRCL with satellite DNA repeats with the identically conserved sequences of the two major piRNAs, but a different arrangement of 21 and 19 alternating repeats (Fig. 6, second row).

**Figure 6.**
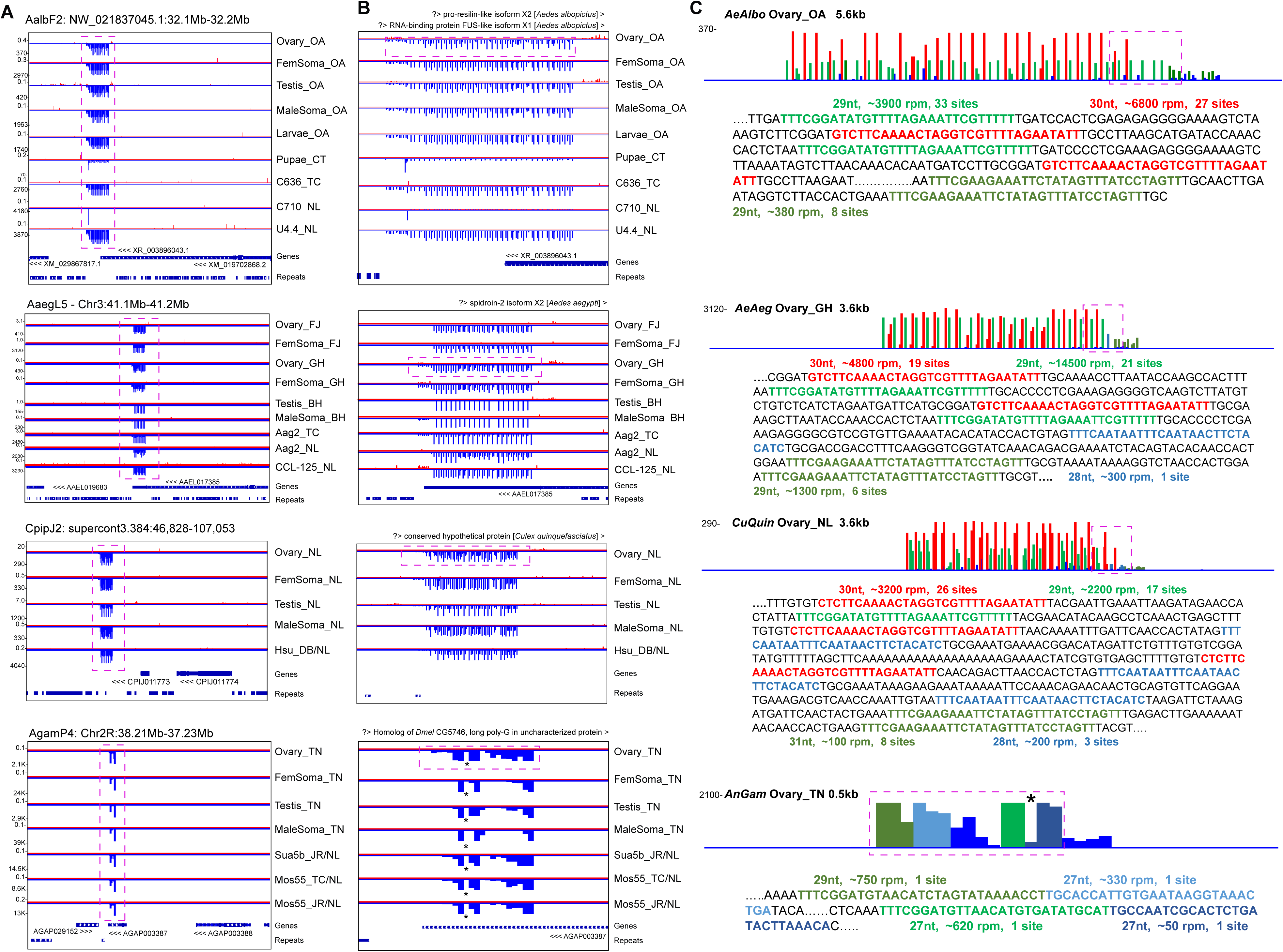
A dynamically evolving Mosquito-Conserved piRNA Cluster locus (MCpiRCL) expressed throughout gonads, soma and cell cultures. (A) Zoomed out genome browser window views at the kilobase level of the MCpiRCL. (B) Zoomed in view of the MCpiRCL from the dashed box in (A). The descriptions of the nearest transcript are listed at the top of the browser window. (C) Microscopic view with reverse complementation of the MCpiRCL from the dashed box in (B). The peaks are color-coded according to the specific reads as DNA in the sequence below each diagram, derived from the region highlighted by the dashed box above the sequence. Reads per million (rpm) and how many occurrences of the read in the satellite tandem repeats within this MCpiRCL. The * in the AnGam MCpiRCL is not accurately representing the read coverage at that spot because of an interval display glitch in the genome browser application.

The *CuQuin* orthologous genic piRCL also displayed these highly expressed satellite DNA repeats where again just two alternating piRNA sequences from 17 and 26 repeats could generate >∼5000 rpm throughout the gonads, somatic tissues, and the Hsu cell line as well (Fig. 6, third row). One of the satellite piRNA’s primary sequence that is 29nt long, “UUUCGGAUAUGUUUUAGAAAUUCGUUUUU”, is perfectly conserved across the ∼100 MYA of evolution (Fig. 1A) but its repeat number has evolved from 17 sites in *Culex* to 21 and 33 sites in *Aedes*. Strikingly, the other *Culex* satellite piRNA sequence only differs from the *Aedes* sequence by just the first nucleotide of 5’-“C” in *Culex* and 5’-“G” in *Aedes,* not only in one repeat site but all 26 repeats in *CuQuin* versus the 19 and 27 sites in *AeAeg* and *AeAlbo*, respectively (Fig. 6C). The most parsimonious explanation for this striking type of sequence evolution suggests a base change first in the culicine ancestor and then parallel evolutionary expansion of the mutated piRNA sequence to form these satellite DNA repeats.

In accordance with the long divergence between culicine and anopheline mosquitoes, *AnGam* appears to lack piRCLs containing satellite DNA repeats, however the orthologous genic piRCL indeed extends to the *AnGam* gene AGAP003387 (Fig. 6, fourth row). In contrast to the culicine genic piRCL, this *AnGam* piRCL is very compact at ∼500bp long within the 3’UTR of AGAP003387 with no tandem repeats but has four main piRNAs comprising >∼1500 rpm. Two of these *AnGam* piRNAs were perfectly conserved at the primary sequence level as one of the culicine satellite DNA piRNAs (Fig. 6C), and this *AnGam* piRCL was also abundantly expressed in *AnGam* gonads and cell cultures. The gene AGAP003387 only has homologs within other mosquitoes, whereas a neighboring gene AGAP003388 is homologous to the *Dmel* gene CG5746 that does generate sone 3’UTR piRNAs (Chirn et al. 2015). Therefore, we have named this region as a Mosquito-Conserved piRNA Cluster Locus (MCpiRCL).

The *AnGam* piRCL may represent the ancestral mosquito locus > ∼200 MYA that began as genic region already primed to express this compact but evolutionarily important piRCL. As the culicine branch evolved greatly expanded genomes from the influx of transposon and other non-coding sequences, the MCpiRCL also expanded to become a satellite DNA repeat perhaps as means to quickly amplify piRNA expression in response to a new selective pressure on its growing genome size. This satellite DNA piRCL has also recently been discovered in *AeAeg* by (Halbach et al. 2020), and has been proposed to trigger the turnover of maternally-deposited transcripts during embryogenesis, similar to the vertebrate tandem repeat cluster of miRNAs miR-430 and miR-427 (Giraldez et al. 2006; Lund et al. 2009). However, whereas miR-430 and miR-427 expression is restricted to the embryo (Giraldez et al. 2006; Lund et al. 2009), the MCpiRCL in all four of these mosquitoes is expressed throughout the gonads, somatic tissues, and cell culture lines (Fig 6B), suggesting the targeting capacity of these piRNAs may be broader than maternally-deposited transcripts.

To predict targets based on a miRNA-targeting configuration, we applied a Smith-Waterman-based algorithm that selected for ‘seed sequence’ matching of bases 1-8 and 2-9 of the piRNA to mosquito gene transcripts, transposons and arboviruses. We predicted many hundreds of transcripts and highlight the top two mRNA, transposon, and virus targets in **Figure S8**. Although the incomplete draft CpipJ2 genome assembly and annotation (Arensburger et al. 2010) may be limiting the number of predicted *CuQuin* targets, there is an expanded repertoire of potential gene and transposon targets for the *AeAeg* and *AeAlbo* piRNAs from this MCpiRCL.

### Culicine mosquitoes exhibit a striking periodicity to the patterns of piRNA biogenesis

Only culicine mosquitoes contained piRCL comprised of satellite DNA repeats (Fig. 6, S4E, S5E, and S7D), and these single highly abundant piRNAs were frequently biased on one strand and were frequently spaced out from each other by a >29nt gap. This piRCL configuration raises the question of how prototypical is the phasing-based mechanism of generating primary piRNAs in mosquitoes versus in *Dmel* where the phased sequential piRNA biogenesis mechanism was first described (Han et al. 2015; Mohn et al. 2015; Pandey et al. 2017; Gainetdinov et al. 2018; Izumi et al. 2020)? We wondered whether culicine mosquito piRNA biogenesis patterns might be distinct from other Dipterans like the anopheline mosquitoes and Drosophilids. Indeed, a previous study applying piRNA phasing algorithms across piRNA datasets from a phylogenetic spectrum of hydra to insects to mammals showed that *AeAeg* piRNAs stood out with the most periodic of 5’ to 5’ piRNA distance peaks (Gainetdinov et al. 2018).

By applying the same algorithm from the previous study (a LOWESS non-parametric regression and auto-correlation smoothing, (Gainetdinov et al. 2018)) to a wide number of *Dmel*, *AnGam*, *CuQuin*, *AeAeg* and *AeAlbo* libraries, we confirmed the strong conservation throughout Dipterans of the one piRNA phasing mechanism that juxtaposes the 3’ terminus of the upstream piRNA to the 5’ start of the downstream piRNA (**Figure 7**). There is also a second phasing pattern where the 5’ starting nucleotide positions of additional downstream piRNAs are rigidly ordered in phase behind the first piRNA, and this periodic 5’-to-5’ phasing pattern is strikingly more evident in the *CuQuin*, *AeAeg*, and *AeAlbo* samples, both in mosquito tissues and in the cell culture lines (Fig. 7). However, this periodic pattern was missing or highly dampened in *AnGam* and *Dmel* samples, with perhaps only *Dmel* ovarian small RNAs subjected to beta-elimination showing the enhanced periodic signal (Song et al. 2014). We hypothesized that perhaps the beta-elimination reaction was clearing some piRNAs not yet fully matured to have 3’-terminal 2’-O-methylation by *Hen1* (Horwich et al. 2007; Kirino and Mourelatos 2007; Saito et al. 2007; Kamminga et al. 2010), thereby enhancing the periodic signal from only fully matured piRNAs. However, we could not increase detection of this 5’-to-5’ periodicity on our other *Dmel* and *AnGam* samples after beta-elimination (Fig. 7 – new samples with “BetaE” label).

**Figure 7.**
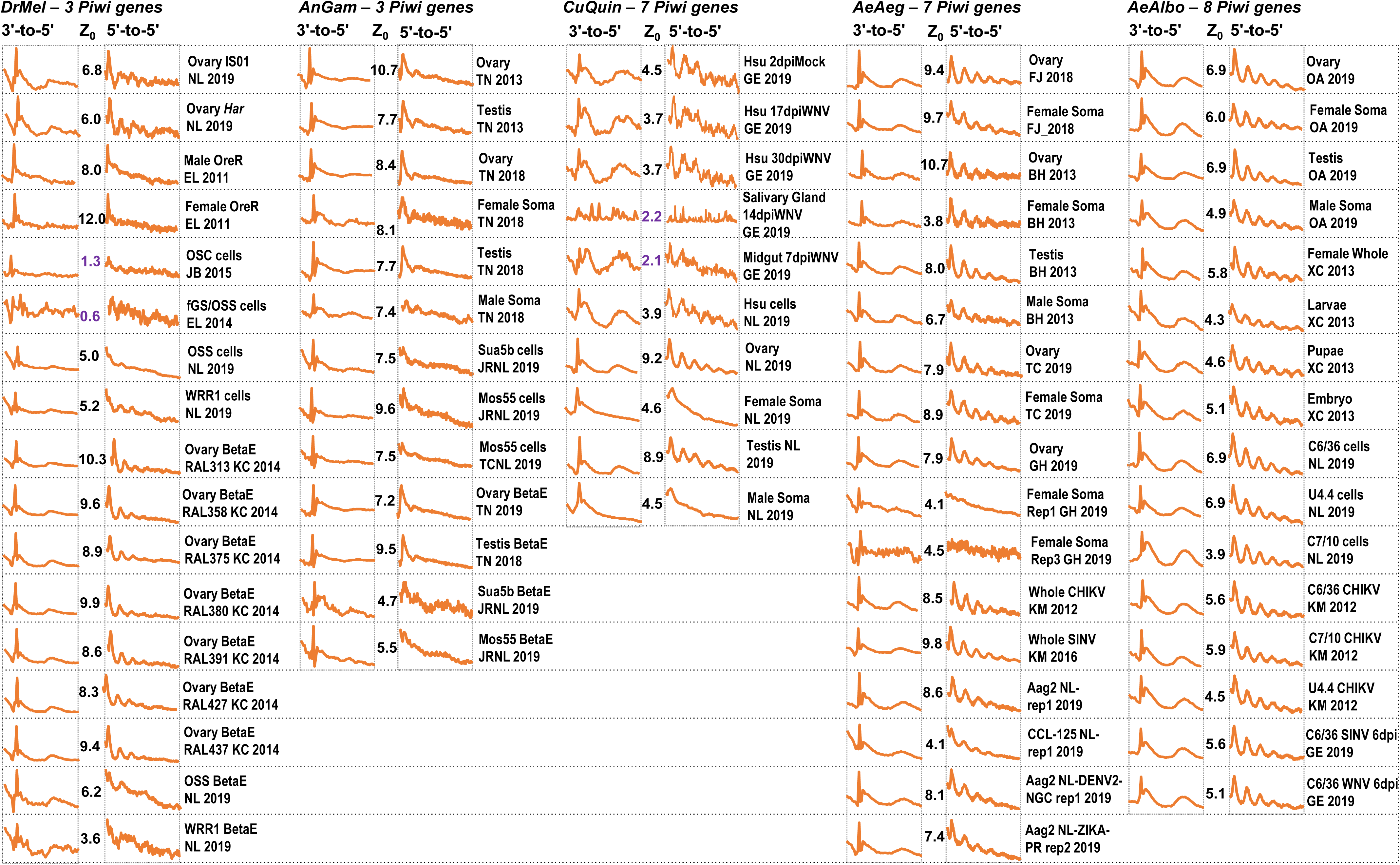
Mosquitoes with expanded Piwi-pathway gene numbers display striking periodicity of 5’-to-5’ piRNA biogenesis phasing patterns. Autocorrelation analysis of piRNAs applied to multiple small RNA libraries from independent labs supports a biological process rather than a technical feature in the detection of this periodic pattern.

Therefore, we speculate that expansion of Piwi pathway genes in culicine mosquitoes may be a factor in promoting greater periodicity in the phasing of piRNA biogenesis patterns and enabling the innovation of satellite DNA repeats in piRCL. To examine the evolutionary relationships of the Piwi pathway genes in Dipterans, we catalogued the *Dmel* genes genetically determined to be in the Piwi pathway, and then conducted blastp and manual curation between NCBI Genbank and VectorBase to arrive at clarified list of Dipteran gene homologs (**Table S7**). We inspected genomic coordinates and blastp outputs to remove several obsolete and redundant gene entries, and we used the blastp similarity scores to match orthologs. Ten core Piwi pathway genes in *Dmel* had single orthologs in *AnGam* that were then frequently expanded into multiple homologs in culicine lineages such as each culicine mosquito having 6 homologs to *Dmel piwi* or *aub* (**Figure S9A**). *AeAlbo* stands out further from *AeAeg* and *CuQuin* by containing the greatest number of expanded Piwi pathway gene families including two *Ago3* homologs and three homologs of *valois* and *vreteno* (Fig. S9A). Another fifteen Piwi pathway genes from *Dmel* had single orthologs in mosquitoes (Fig. S9B). Indeed, the expansion of *piwi* and *aub* homologs in culicine mosquitoes is likely to explain piRCL innovation such as *AeAeg* PIWI4 appears to be most critical for loading the satellite repeat MCpiRCL (Halbach et al. 2020). Although seven *Dmel* genes including well known factors in *Drosophila’s* piRNA-mediated transcriptional gene silencing mechanisms (i.e. *panx, rhi, del,* and *cuff* (Le Thomas et al. 2014; Mohn et al. 2014; Zhang et al. 2014)) were completely absent in mosquito genomes, this may foretell potential mosquito-specific factors required for its unique repertoire of Piwi pathway genes.

## DISCUSSION

Cell culture lines are invaluable resources for genomic and biochemical studies that complement genetic studies in the organism. This feature is clearly demonstrated by the important banks of genomic, transcriptomic and epigenetic datasets for model organism and human cell lines in the ModENCODE and ENCODE projects, respectively (Graveley et al. 2011; Kharchenko et al. 2011; Negre et al. 2011; Consortium 2012; Djebali et al. 2012; Thurman et al. 2012). Mosquito cell cultures from various species (Fig. 1C) also serve such a vital role in arbovirus research by facilitating virology and biochemical studies, but we are only now beginning to better characterize these cell lines for comparing to the tissues of the animal. In this study, we bring together an integrated resource of small RNA genomics analyses for four medically relevant mosquito species with genome annotations that are furthest developed by the arbovirology and entomology communities.

Mosquitoes arguably may have a more translational impact on human health compared to model organisms studied in the ModENCODE project, yet genomic characterizations of the culicine mosquitoes in particular have lagged because their significantly larger genomes are inflated by a large proportion of transposons and other repetitive elements. New genomic approaches such as high-throughput long-read and Hi-C sequencing approaches have only recently bridged scaffolding gaps to bring about major improvements in the *AeAeg* and *AeAlbo* genome assemblies (Dudchenko et al. 2017; Matthews et al. 2018; Palatini et al. 2020). However, functional annotations such as improving gene models with better transcriptome data is still needed for mosquito genomics advancement including this study in which we opted to analyze the CpipJ2 assembly that had genes and repeats tables (Arensburger et al. 2010) but was still much more fragmented than the newer CpipJ3 assembly which lacked annotation (Dudchenko et al. 2017). Our study also demonstrates the need for better repetitive elements annotations including refinement of transposons beyond the automated programs like RepeatModeler (Wheeler et al. 2013; Flynn et al. 2019) which generate comprehensive but redundant repeats list. Strangely, the majority of mosquito piRNAs across species do not appear to target transposons and may ultimately have a wide range of other targets yet to be determined.

As the diversity of *Dmel* cell culture lines has greatly expanded just in the last decade, only 4 *Dmel* lines are known to express piRNAs (fGS/OSS, OSS-OSCs-OSC-delta-MBT, WRR1 and Kc cells, (Lau et al. 2009; Saito et al. 2009; Fagegaltier et al. 2016; Sumiyoshi et al. 2016; Vrettos et al. 2017)), while the vast majority of *Dmel* cell lines only express miRNAs and siRNAs (Wen et al. 2014). Such few lines may reflect the exceptional nature of *Dmel* to restrict Piwi pathway gene expression just to the gonads, whereas most other insects robustly express piRNAs in the soma (Lewis et al. 2018). The smaller selection of mosquito cell cultures were generated many decades ago from very heterogenous tissue sources (Table S1) coupled with the long term passaging and wide distribution of these cells have likely selected for cells with gene expression profiles that are extremely diverged from specific mosquito tissues. Every mosquito cell line in this study expressed piRNAs, including our culture of C7/10 cells (Fig. 5B) that may differ from a previous report of C7/10 cells that lacked piRNAs (Skalsky et al. 2010); yet some mosquito lines abundantly expressed somatic piRNAs while other isolates were quite subdued in somatic piRNA expression. With this initial sampling of cell cultures and wild-caught versus domesticated lab isolates, the data suggests that somatic piRNAs and siRNAs may be an insect vector response to a persistent arbovirus infection. We envision a future effort to profile geographically-diverse set of wild mosquito isolates that we will add to the Mosquito Small RNA Genomics Resource (MsRG, http://class.bu.edu/~nclau/MSRG_db/MSRG_db_Home.html).

In addition, the MSRG will enable future virology and biochemistry experiments with mosquito cell cultures to be placed in better context to experiments with mosquitoes. For example, our hierarchical clustering of miRNA and transposon small RNA profiles show that cell cultures have a transcriptome state that is distinct from gonads and somatic tissues (Fig. S3, S4, S5, S7), and these results are further supported by additional Principal Component Analysis (PCA) plots (**Figure S10**). However, the PCA plots also suggest that different labs’ isolates of *AnGam*, *CuQuin* and *AeAeg* cell cultures showed a higher degree of clustering together than the cell lines from *AeAlbo*. Finally, the MSRG provides a reference list of curated mosquitoes genic and intergenic piRCLs (Fig. S10C) and reference lists of mosquito arboviruses and transposons with abundant small RNAs from both cell cultures and colonies. This resource provides the necessary foundation to accelerate and better connect all the different mosquito RNAi pathways with arboviral research within an important vector organism.

## MATERIALS AND METHODS

### Mosquito strains, cell cultures and virus infections

The *AnGam* isolate from Imperial College, UK was kept in standard rearing conditions as in (Castellano et al. 2015). The *AeAeg* isolates from Colpitts lab were maintained in the insectary of the National Emerging Infectious Disease Laboratory (NEIDL) as described in (Araujo et al. 2020). The *AeAeg* isolate from the Hughes lab were maintained in the insectary at the University of Texas Medical Branch as described in (Saldana et al. 2017). The *AeAlbo* isolates from the Akbari lab were described in (Gamez et al. 2020). The *CuQuin* isolates were purchased from Benzon Research.

All mosquito cell culture media are described in Table S1, and all cultures were established in the Lau to grow stably for months before cells were used for total RNA extraction and multiple live aliquots were cryopreserved. Cells were all kind gifts: Sua5b and Mos55 cells from the Rasgon lab; C6/36 and Mos55 cells from the Colpitts lab; Aag2 cells from the Blair lab, CCL-125 from the Connor lab; C7/10 cells from the Fallon lab; and U4.4 and Hsu cells from the Brackney lab. All cells were maintained in a humidified incubator at 28C with 5% CO2 atmosphere. The DENV and ZIKV infections were performed on Aag2 cells that were ∼80% confluent in T25 flasks grown in Shield & Sang Media (Table S1) using viral supernatants from previous C6/36 infections. The infections were conducted in the BSL2+ facility in the NEIDL and were cultured for 7 days before cells were neutralized in the TRI-reagent for total RNA extraction. Viral infection status was confirmed by the qRT-PCR assay detailed in (Araujo et al. 2020).

### Small RNA library preparation and deep sequencing

Small RNA libraries were constructed either from small RNAs size fractionated from Urea-Polyacrylamide Gel Electrophoresis as in (Chirn et al. 2015) or enriched for from Q-sepharose matrix as in (Srivastav et al. 2019). For size-fractionation of small RNAs, 1-5 ug of total RNA from mosquito tissues and ∼10ug of total RNA from cell lines was extracted with TRI-reagent. Size fractionation was performed on a urea-denaturing 15% polyacrylamide gel with TBE buffer and 18-nt and 32-nt fluorescent oligos were used as markers. 18-32nt sized RNA portion of gel was excised under UV and eluted in 500µL 0.3M NaCl overnight with mild agitation at RT. Small RNA containing eluate was saved and supplemented with 2 volumes of ethanol and 1µL of 20mg/mL glycogen for precipitation at −20°C overnight. Small RNAs were precipitated by centrifuging at 15,000rpm at 4°C for 20 mins. Small RNA containing glycogen pellet was next washed with chilled 75% ethanol and eluted in 12µL of freshly made 50% (w/v) PEG-8000 to enhance 3’ end ligation efficiency. 6µL of the small RNAs in PEG-8000 were used for library construction using NEBNext Small RNA Library Construction kit (E7330S) as per manufacturer’s protocol.

For Q-sepharose enrichment of small RNAs from cell lines, 25-50µL of pelleted cells were eluted in 500µL of binding buffer (20 mM Hepes pH 7.9 (with KOH), 10% glycerol, 100 mM KOAc, 0.2 mM EDTA, 1.5 mM MgCl2, 1.0 mM DTT, 1X Roche Complete EDTA-free Protease Inhibitor Cocktail). Cells were lysed using one freeze-thaw cycle and pulverizing with a plastic pestle. Separately 1.5ml aliquot of Q-sepharose FF matrix suspension was washed 1X in water, then 3X in binding buffer, then mixed with the cell lysate at mild agitation at 4°C. Next, 300mM KOAc was added to the suspension. For *Drosophila* cells, at this step 2S rRNA gets bound by the Q-sepharose, while small RNA RNPs remains in the supernatant. Next, the Q-sepharose suspension was spun at 2000 rpm for 5 mins to isolate all the small RNA RNPs in supernatant followed by iso-propanol small RNA extraction from the supernatants. Small RNAs extracted were also eluted in 50% (w/v) PEG-8000 and library constructions similarly as above.

All small RNA libraries were purified with the Monarch PCR & DNA Cleanup Kit (5 μg), quantified using Qubit 2.0 and analyzed on Agilent Bioanalyzer 2100 before sequencing on the BUSM Microarray and Sequencing Resource. For total RNA from *Drosophila* OSS and WRR1 cells and AnGam Sua5b and Mos55 cells, we subjected this to beta-eliminition treatment as in (Song et al. 2014).

### RT-PCR analysis of *AnGam densovirus* in Mos55 cells

Total RNA was extracted from Mos55 cells by TRI-reagent RT, and 10ug RNA was subjected to DNaseI and RNAse A digestion for 30 minutes at 37°C, heat-inactivated at 65°C, and then subjected to standard phenol-chloroform:IAA extraction and isopropanol precipitation. First strand cDNA synthesis was performed using ∼1.0 µg untreated RNA, 0.78 µg DNaseI-treated RNA, and 0.25 µg RNaseA-treated RNA using the NEB Random Primer Mix and Protoscript. PCR was performed on 1uL of Mos55 cDNA in 50 µL reactions using the specified Amp1, Amp2, and *AnGam* Rps7 primer pairs with Phusion High-Fidelity DNA Polymerase. Amp1 primers: TACAAGAACAAGGCAGTTCCAGC; CCAATAAGTTATCCAATATTAGTG. Amp2 primers: TGGACTTATATCAAATTCCTATATGG; ACGGGGATCCCGGACTAATGTTGGC. *AnGam* Rps7 primers: GGTGCACCTGGATAAGAACCA; CGGCCAGTCAGCTTCTTGTAC.

### Reducing redundancy in transposon family consensus sequences lists

Since most mosquito transposon annotations have been derived automatically with bioinformatic prediction scripts such as the RepeatModeler package that consists of RepeatMasker, RepeatScout/TEFam, RECON and TRF program tools (Bao and Eddy 2002; Price et al. 2005; Gelfand et al. 2007; Wheeler et al. 2013), the RepeatModeler’s heuristic issue is that its efficient process generates lists of transposon families that are very redundant. Therefore, we developed different strategies for each specie to mitigate over-counting of small RNAs.

From the 3607 *AnGam* transposon sequences we retrieved from VectorBase (Giraldo-Calderon et al. 2015), we first removed all sequences from non-*AnGam* organisms. We then grouped the remaining sequences into Repbase (Kapitonov and Jurka 2008), NCBI, EMBL, TEFam (Wheeler et al. 2013) categories based on the sources of the transposon sequences. Within the Repbase group, we manually attached the LTR segments to the associated internal regions; then we performed an all-against-all pairwise search using BLAT with 90% sequence identity cutoffs (Kent 2002). Beginning with the cluster centroids from Repbase, we progressively added sequences from NCBI, EMBL, and TEFam into the existing clusters. If no existing centroid has over 90% sequence identity with the current sequence, the current sequence is designated as the new centroid. In the end, we consolidated our list to 555 *AnGam* transposon consensus family sequences.

From the 4817 *AeAeg* transposon sequences we retrieved from VectorBase, we removed all sequences from non-*AeAeg* organisms; we then grouped the remaining sequences into Repbase, TEFam, NCBI, Vectorbase, Recon categories based on the sources of the transposon sequences. Within the Repbase group, we again manually attached the LTR segments to the associated internal regions. We use MeShClust version 1 (James et al. 2018) with 55% sequence identity cutoff to reduce redundancy within each source of Repbase, TEFam, NCBI, Vectorbase, and Recon. Then we performed an all-against-all pairwise search using BLAT with 55% sequence identity cutoffs. Beginning with the cluster centroids from Repbase, we progressively add sequences from TEFam, NCBI, VectorBase, and Recon into the existing clusters. If no existing centroid has over 55% sequence identity with the current sequence, the current sequence is designated as the new centroid. This resulted in 1242 *AeAeg* transposon consensus family sequences.

For the 3730 *CuQuin* transposon sequences we retrieved from VectorBase that were automatically identified by RepeatModeler version 1, we first grouped them into 59 families based on sequence annotations, e.g., LTR_Gypsy. Then within each group, we used MeShClust version 1 with 55% sequence identity cutoff to reduce redundancy and designated the transposon sequence with highest average read counts across several small RNA datasets as the cluster centroid. This resulted in 751 *CuQuin* transposon consensus family sequences.

For *AeAlbo*, we first started with 8556 transposon sequences automatically identified by RepeatModeler version 2 (Flynn et al. 2019) from the AalbF2 assembly (Palatini et al. 2020), that we grouped into 66 families based on sequence annotations, e.g., LTR_Gypsy. Within each family, we used CD-HIT with 55% sequence identity cutoff to cluster transposon sequences, designating the transposon sequence with highest average read counts across several small RNA datasets as the cluster centroid. We then removed transposon sequences that were less than 500 nucleotides as result of the higher LTR-detection sensitivity of RepeatModeler version 2 (Flynn et al. 2019). This resulted in 3098 *AeAlbo* transposon consensus family sequences.

From these consolidated lists, we applied the RepeatMasker program (Wheeler et al. 2013) to identify the genome copy numbers and genome coverages for each transposon from four organism, and then applied small RNA counts for the benchmarking results in Fig. S1.

### Bioinformatics analysis of small RNA datasets

For these mosquito species, we adapted our bioinformatics analysis pipelines for analyzing genic/intergenic small RNA counts and analyzing transposons/virus counts (Chirn et al. 2015). We first indexed the genome assembly file by running BWA version 1 (Li and Durbin 2010) and formatdb from NCBI. Within the genic/intergenic small RNA pipeline, small RNA reads were first trimmed by Cutadapt program (Didion et al. 2017) to remove the adaptor sequences in the 3’ end. Trimmed reads were then mapped to a collection of virus sequences using Bowtie1 with 2 mismatches (Langmead et al. 2009). Reads which were mapped to the virus were removed. Next, reads were mapped to miRNAs and structure RNAs, e.g. snRNAs, tRNAs, rRNAs, snoRNAs using Bowtie1 with 2 mismatches. Reads which were mapped to miRNAs and structure RNAs were removed. Finally, reads were mapped to genomes using Bowtie1 with 2 mismatches to get the genic/intergenic counts using the genome GTF file. Genic counts were further categorized into 5’UTR counts, CDS counts, 3’UTR counts.

The fixed step Wig file was generated by recording the normalized read counts within every window of 25 bases for positive strand and negative strand respectively. The wigToBigWig program was used to covert the fixed step wig file to the bigwig file which was loaded to the Broad Institute Integrative Genomics Viewer (IGV(Robinson et al. 2011)) together with the genome assembly and GTF files. Reads mapped to the intergenic regions were progressively clustered together if normalized read counts is over 0.02 within a sliding window of 25 base. To reduce the redundancy in the genic table caused by different isoforms of a gene, the mergeBed program (Quinlan 2014) was used to consolidate different isoforms by providing the genomic location of each isoform. The isoform with the highest read counts was chosen as the representative of the gene.

Within transposons/virus sRNA pipeline, reads were first trimmed by Cutadapt program to remove the adaptor sequences in the 3’’ end. Then trimmed reads were mapped to miRNAs with BLAST (Altschul et al. 1990). Reads which were mapped to miRNAs were removed. Then reads were mapped to transposons using Bowtie1 with two mismatches and virus using Bowtie1 with one mismatch. Finally, the mapping patterns with respect to transposons/viruses were plotted with an R script. Hierarchical Clustering was performed by calling Python Seaborn Clustermap function using Euclidean distance and average linkage clustering method. Principal Component Analysis (PCA) was carried out by R prcomp function, with plots generated by the ggplot function.

### Curation of genic and intergenic piRNA Cluster Loci (piRCL)

Tables for genic and intergenic piRNA clusters were manually curated to remove piRNA calls that may be inaccurate due to the incompleteness of a genome assembly. To curate genic piRCL, we filtered out piRNA cluster calls with very low expression and in genes with usual structures (extremely long introns and very small exons). First, consolidated tables were sorted to remove piRNA entries that were in genes with lengths >100Kb. Then, entries with normalized locus RPM values <10 were removed. Read concentration values, calculated as (Normalized locus/Length)*1,000, were calculated for the resultant entries and then sorted from highest to lowest. Entries with read concentration values <15 were removed. Finally, each entry was manually checked in the IGV browser to remove entries with either one or both of the following characteristics: IGV window y-scale values <10 and genes with extremely large introns >10Kb.

Intergenic piRCLs were manually curated by first ordering piRNA loci entries and scanning RPM(all) values >100 and >1,000. Then, we searched among the ordered entries to find highlighted RPM(all) values to narrow our searches. Whenever three or more highlighted loci entries were found close proximity, this genomic range was noted and then searched in the IGV browser to visualize the piRNA clusters in this region. In addition, we searched past the left and right bounds of the noted loci to see if more piRNA clusters were present. If additional piRNA clusters were present, the intergenic range was noted. Intergenic cluster regions >1,000bp were kept in a final table and anything smaller was excluded.

After the first level of manual curation, we combined the selected curated loci from all the samples within each species by prioritizing all the loci called in ovaries with SQL commands that added back the other loci absent from the curated ovaries list. The normalized the gene counts were extracted from the consolidated genic table for each curated genic piRNA cluster. The mergeBed program was used to sum all the intergenic peaks within a curated intergenic piRNA cluster loci.

### piRNA target prediction from the mosquito conserved piRCL

We extracted the 5’-1 to 8 or 2 to 9 nucleotides of the piRNA sequences as 8mer seed sequences and used the Water program which implements Smith-Waterman search algorithm (Smith and Waterman 1981) to look for the perfect match against the reverse strands of transcripts, transposons, and viruses. Once the perfect 8mer match against the reverse strands was found, we extend hits so that they have the same length as the piRNA sequence. We designated the extended hits as candidates. Then we used Water program to search the full piRNA sequences against the reverse strand of the candidates to identify the total number of perfect matches. In the end, candidates were sorted in the decreasing order of numbers of perfect matches.

### piRNA Ping-pong and Phasing analysis

Reads were first trimmed by Cutadapt program to remove the adaptor sequences in the 3’ end. Then, trimmed reads longer than 23 nucleotides were aligned to the genome using Bowtie 1 with no mismatch. The genomic location and the number of times of mapped reads were recorded. Using this information, we carried out autocorrelation analysis to identify periodic peaks based on a previous script from (Gainetdinov et al. 2018). For Ping-pong analysis, autocorrelation analysis of 5’ to 5’ distance on the opposite genomic strands were carried out and Z score at distance 10 was calculated, a Z score over 2 was observed in most cases. For 3’ to 5’ phasing analysis, autocorrelation analysis of 3’ to 5’ distance on the same genomic strands were carried out and Z score at distance 0 was calculated, a Z score over 2 was observed in most cases. For 5’ to 5’ phasing analysis, autocorrelation analysis of 5’ to 5’ distance on the same genomic strands were carried out and periodic peaks were observed on the autocorrelation scores.

## ACKNOWLEDGEMENTS

We thank Ildar Gainetdinov and Phil Zamore for assistance with the piRNA phasing code, Jullien Flynn for analyzing the *AeAlbo* genome with RepeatModeler v2; the BU Microarray and Sequencing core for timely sequencing; Dianne Schwarz and Mohsan Saeed for comments on the manuscript. SPS, SG, FF, TC, EIP, GH, TN, AG, and EM conducted the experiments; RJ, JR, JHC and DEB provided mosquito cultures, QM developed and executed the bulk of the bioinformatics analyses, and SG also provided extensive help with analysis and curation. NCL conceived and conducted experiments and wrote the paper with comments from all authors. This work was supported by NIH grants R01-AG052465 and R21-HD088792 to NCL; NIH grants R01-AI128201, R01-AI116636, and R01-AI150251 to JLR; Defense Advanced Research Project Agency (DARPA) Safe Genes Program Grant (HR0011-17-2-0047) and an NIH New Innovator Award (1DP2AI152071-01) to OSA; NEIDL Pilot funds and NIH grant R21AI129881 to TMC; NIH grants R21NS101151 and R21AI121933-01 to JHC; the LSTM Director’s Catalyst Fund award to EIP; NIH grants R21AI129507 and R21AI138074), the BBSRC (BB/T001240/1), and a Royal Society Wolfson Fellowship (RSWF\R1\180013) to GLH; grants from BBSRC Network Grant “ANTI-VeC” (AV/PP0020/1) and the Bill & Melinda Gates Foundation to TN. Small RNA sequencing data is deposited in the NCBI GEO as Study #SRP178563 and GSE146545.

## SUPPLEMENTARY TEXT

### Reducing redundancy in transposon family consensus lists for CuQuin, AeAeg and AeAlbo

We could not apply BLAT to *CuQuin*, *AeAeg*, and *AeAlbo* repeats lists because there were too few curated RepBase entries to reliably prioritize the centroid choices, and BLAT was too computationally greedy to handle the RepeatModeler-version1 outputs (Wheeler et al. 2013). Therefore, we applied the MeShClust program as a more computationally-efficient method to cluster redundant repeats entries at a 55% similarity cutoff (James et al. 2018), and we selected the centroid entry prioritized by the rubric of “RepBase > TEFam > NCBI > VectorBase > Recon” (Fig. S1B). We arrived at the 55% similarity cutoff by benchmarking different cutoffs applied to the AeAegL5 genome assembly to see the *AeAeg* repeats list reduced by ∼75% to 1,242 transposon families with optimized small RNA mapping and importantly still preserving the ∼60% genomic coverage. We also applied this method to the *CuQuin* repeats families by reducing the transposon consensus sequences and small RNA double-counting by 46% and 34% respectively (Fig. S1C).

However, we had to further modify our transposon family reduction approach for the latest AalbF2 assembly (Palatini et al. 2020) because the original transposon family consensus sequence list was derived from an *AeAlbo* C6/36 cell line rather than de novo from the AalbF2 assembly (Thibaud-Nissen, F. personal communication). Therefore, we applied to this assembly the RepeatModeler-version 2 (Flynn et al. 2019) which has a more sensitive Long Terminal Repeats (LTRs) detection module, and we generated a first list of 8,556 repeats families of which 5,020 of these were LTRs. This high number of entries was clearly redundant because this covered an excessive 85% of the genome (Fig. S1D) yet was also too large an input file for MeShClust to handle. We overcame this problem by then using the CD-HIT program (Li and Godzik 2006) for clustering centroids from the AalbF2 assembly’s RepeatModeler_v2 outputs and arrived at just 3,098 transposon families covering ∼65% of the genome, thereby being more in line with the genomic coverage percentage of *<u>AeAeg</u>* transposons.

### Additional details on how we assessed viral infections of Aag2 cells

To T25 flasks of ∼80% confluent Aag2 cells cultured in Shield and Sang M3 media and 10% FBS, we applied C6/36 cell supernatants each previously subjected to an extensive arbovirus infection. Our first experiment achieved decent viral infection levels in Aag2 cells, but were enhanced 2 orders of magnitude more in the second infection experiment, which were still Aag2 cells cultured in the Lau lab but from a longer passage than the first batch of cells (Fig. S6C). Although there were greater proportional levels of viral siRNAs and piRNAs in experiment 2 (Fig. S6D, S6E), the small RNA increases were not directly proportional to the differences in the viral RNAs, and the Mock Control from this second batch of Aag2 cells already exhibited higher levels of inherent virus such as PCLV, CFAV, CYV, and Densovirus. Perhaps the longer passaging of the second batch of Aag2 cells built up a larger reservoir of RNAi factors that also allowed higher tolerance of greater virus replication.

## SUPPLEMENTARY FIGURE LEGENDS AND TABLE LEGENDS

**Figure S1.**
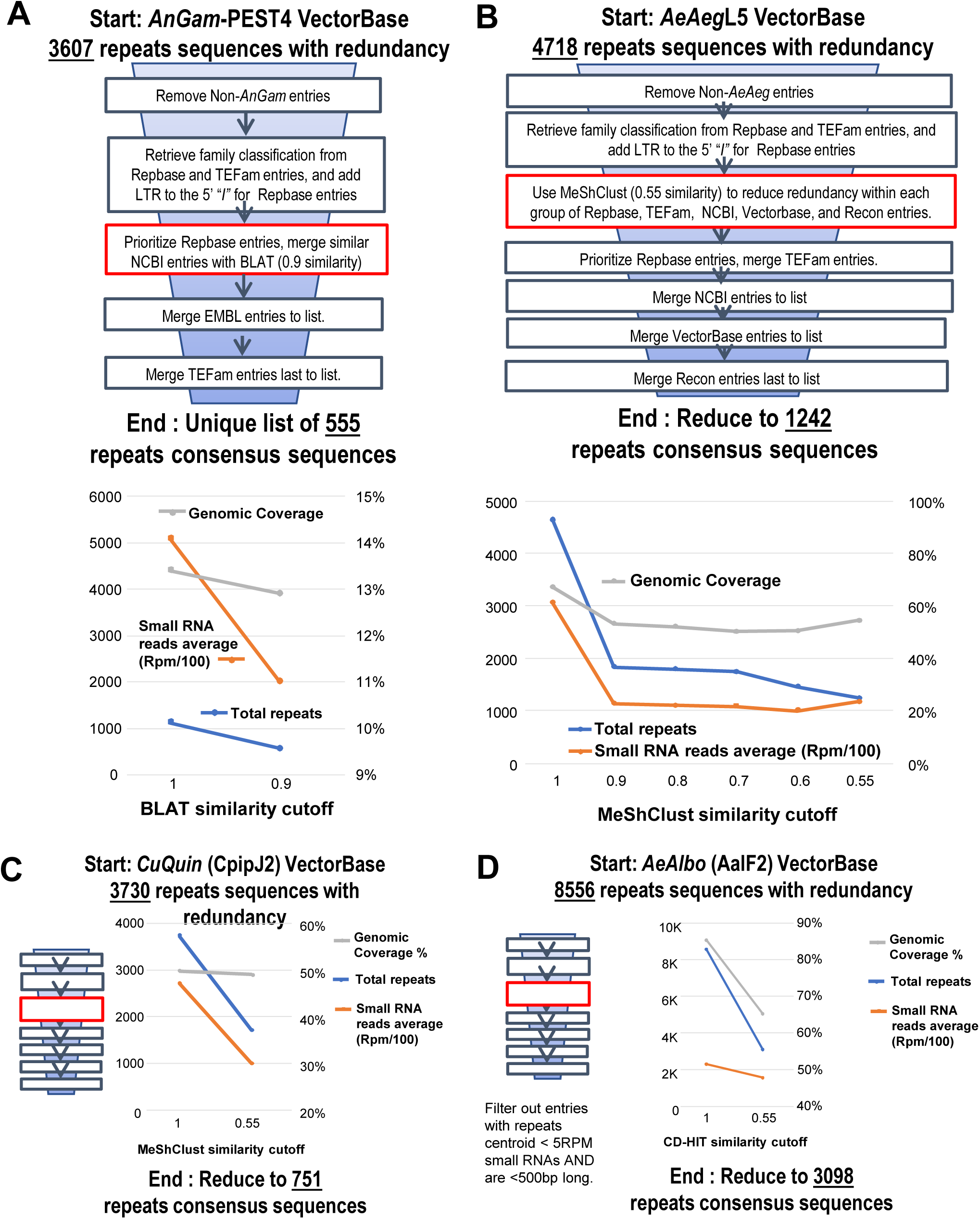
Methodology to reduce redundancy in mosquito transposon family annotation lists. (A) Top: Diagram of the pipeline to condense the *AnGam* repeats families list, with critical parameters for the BLAT algorithm in the red box. Bottom: Benchmarking results from reducing the *AnGam* repeats list to 555 repeats families. (B) Top: Diagram of the pipeline to condense *AeAeg* repeats families list, with the critical parameters for the MeShClust algorithm in the red cox. Bottom: Benchmarking results for reducing the *AeAeg* repeats list to 1,242 repeats families. (C) and (D) The ideogram represents the similar pipeline as the *AeAeg* pipeline and benchmarking results from reducing repeats families in *CuQuin* and *AeAlbo* genomes.

**Figure S2.**
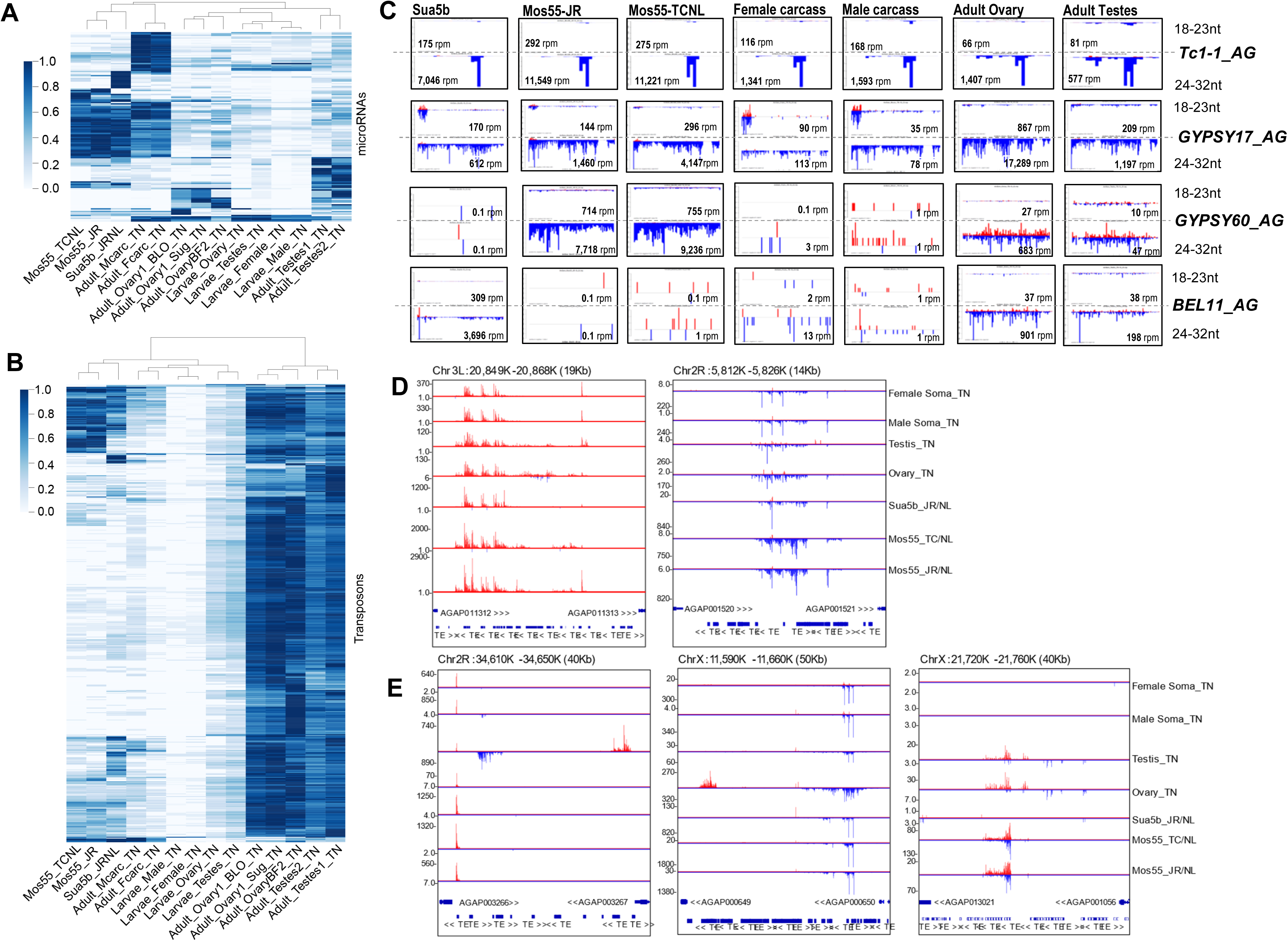
Hierarchical clustering heatmaps of *AnGam* microRNA and transposon small RNA profiles, highly expressed transposon small RNA patterns common to cell lines and in the mosquito; and notable *AnGam* piRNA clusters. Hierarchical clustering of (A) miRNAs and (B) transposon-mapping small RNA patterns for the cell lines versus the mosquito tissues. (C) Profiles of the *AnGam* transposons with most abundant small RNAs both in cell cultures and *AnGam* tissues. (D) Genome browser views of notable genic piRNA Cluster Loci (piRCL) in *AnGam*. (D) Genome browser views of notable intergenic piRCLs in *AnGam*.

**Figure S3.**
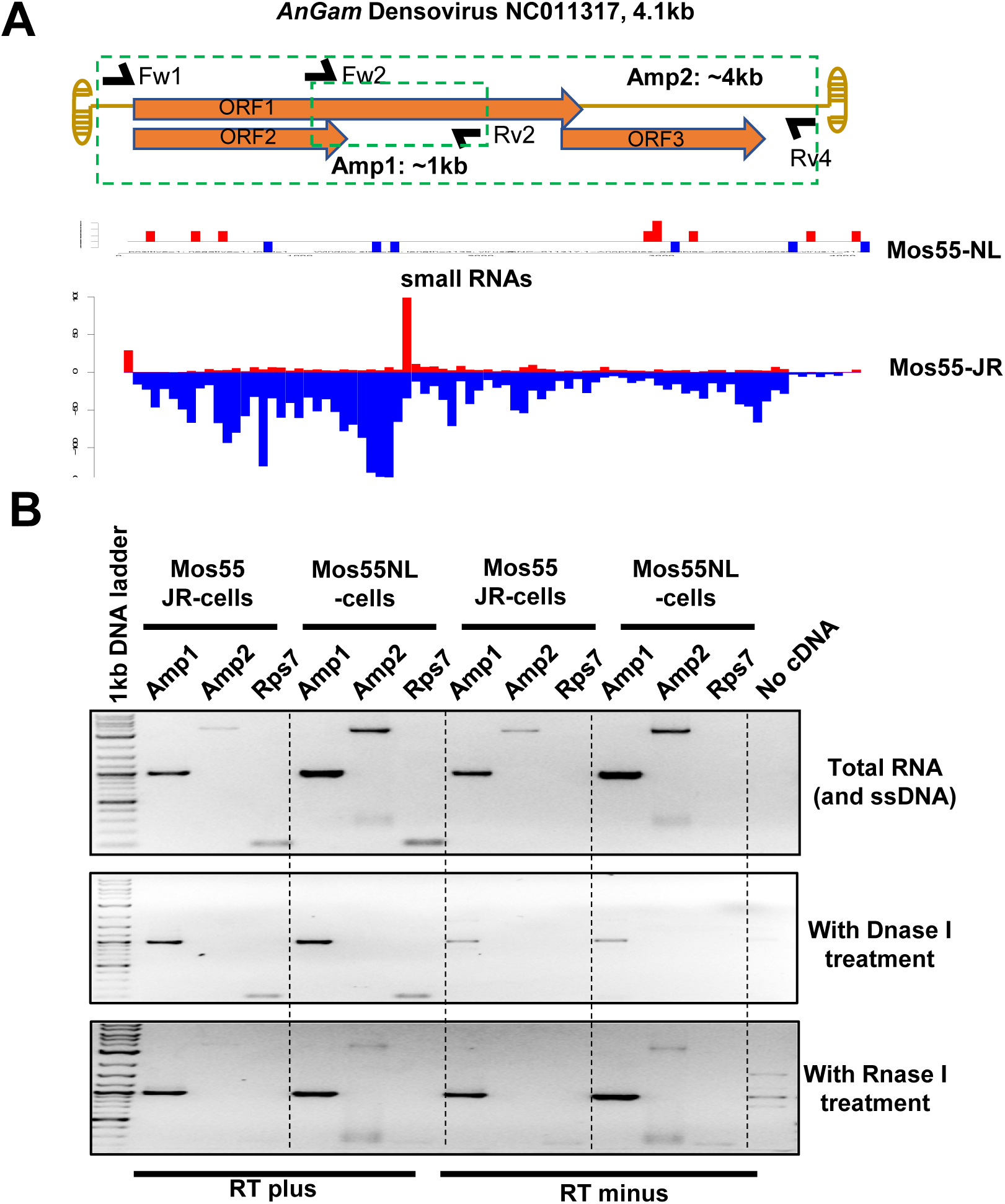
*AnGam densovirus* infection status of Mos55 cells. (A) Schematic of primers used to detect DNA and RNA copies of *AnGam densovirus* NC011317, and the diagrams of viral small RNAs mapping to *densovirus* between Mos55-NL and Mos55-JR cells, which are independent isolates from the Lau and Rasgon labs, respectively. (B) RT-PCR results of amplicons from input of total RNA from Mos55-NL and Mos55-JR cells, with also further treatments with DNase I or RNase I. Amp1 and Amp2 are noted in (A), while Rps7 is a ribosomal protein 7 housekeeping gene as a control mRNA.

**Figure S4.**
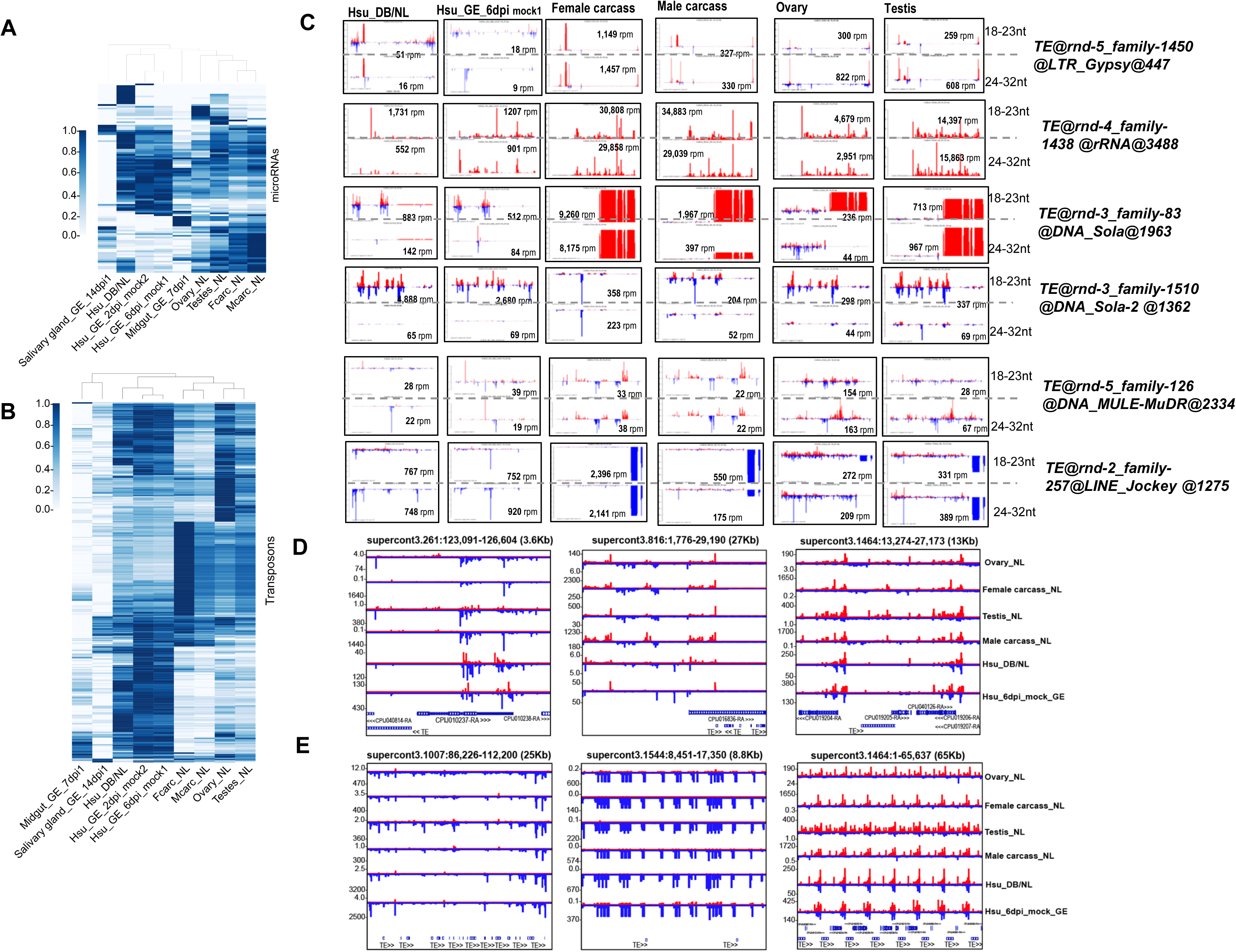
Hierarchical clustering heatmaps of *CuQuin* microRNA and transposon small RNA profiles, highly expressed transposon small RNA patterns common to cell lines and in the mosquito; and notable *CuQuin* piRNA clusters. Hierarchical clustering of (A) miRNAs and (B) transposon-mapping small RNA patterns for the cell lines versus the mosquito tissues. (C) Profiles of the *CuQuin* transposons with most abundant small RNAs both in cell cultures and *CuQuin* tissues. (D) Genome browser views of notable genic piRNA Cluster Loci (piRCL) in CuQuin. (D) Genome browser views of notable intergenic piRCLs in *CuQuin*.

**Figure S5.**
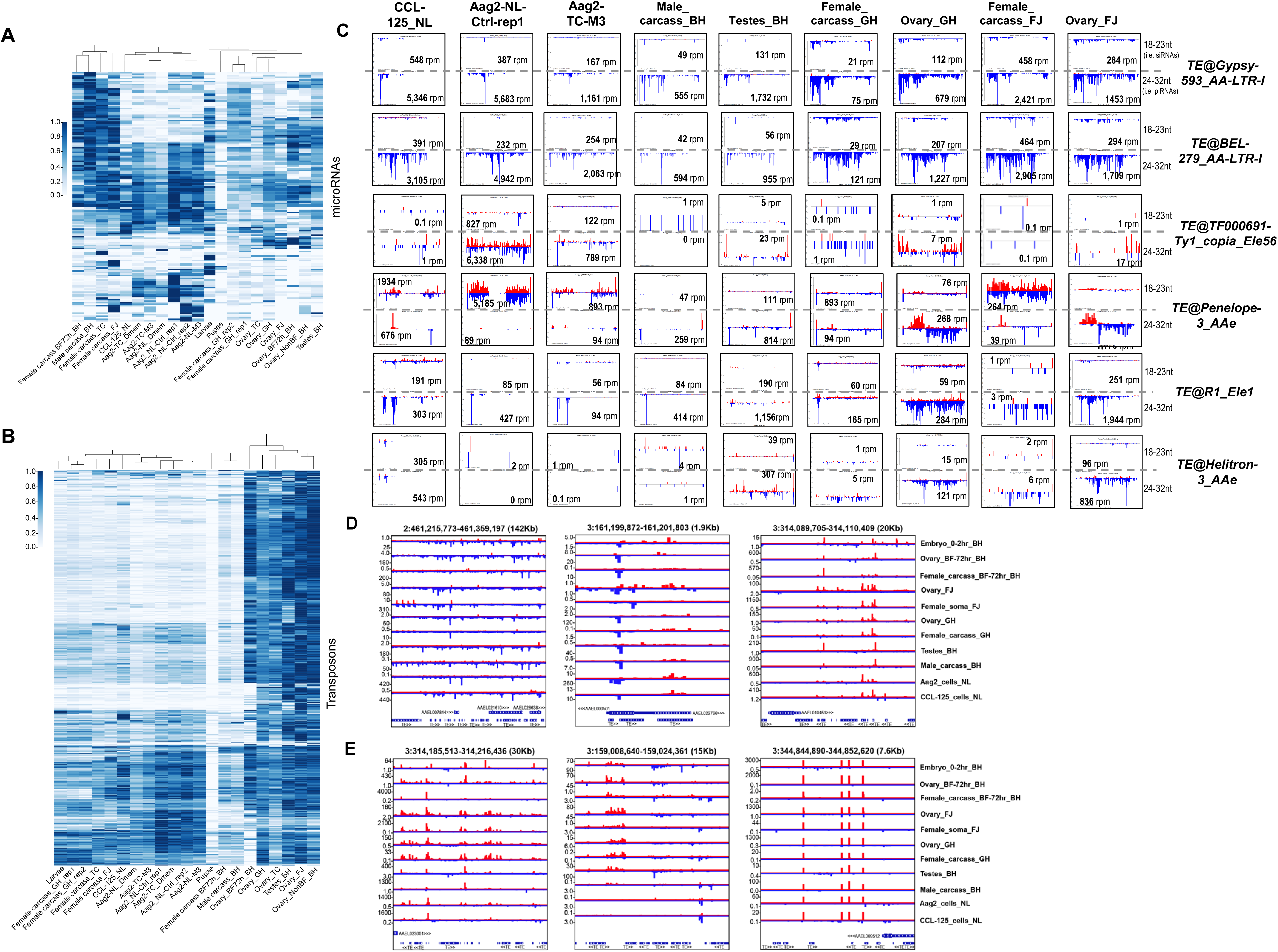
Hierarchical clustering heatmaps of *AeAeg* microRNA and transposon small RNA profiles, highly expressed transposon small RNA patterns common to cell lines and in the mosquito; and notable *AeAeg* piRNA clusters. Hierarchical clustering of (A) miRNAs and (B) transposon-mapping small RNA patterns for the cell lines versus the mosquito tissues. (C) Profiles of the *AeAeg* transposons with most abundant small RNAs both in cell cultures and *AeAeg* tissues. (D) Genome browser views of notable genic piRNA Cluster Loci (piRCL) in *AeAeg*. (D) Genome browser views of notable intergenic piRCLs in *AeAeg*.

**Figure S6.**
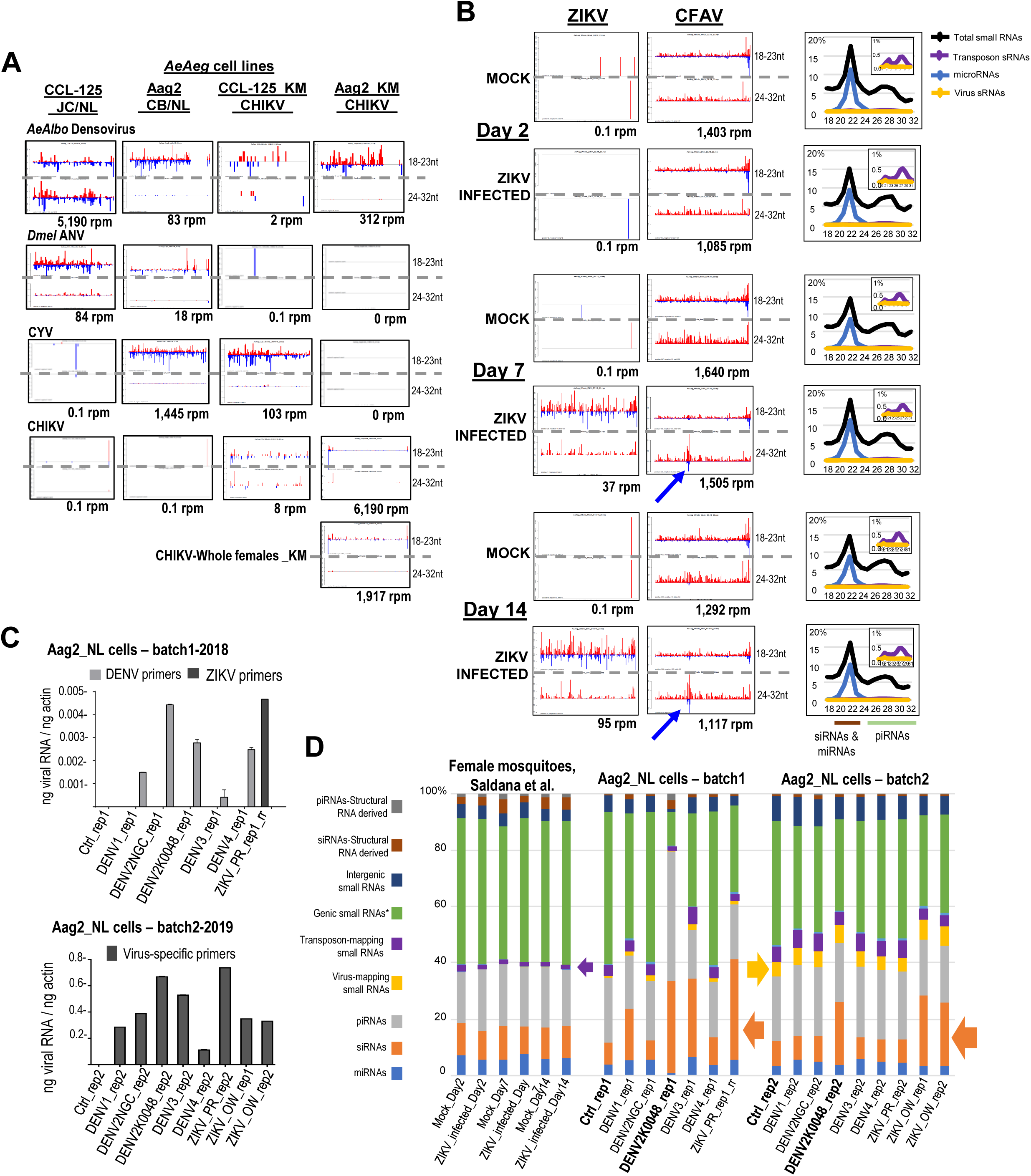
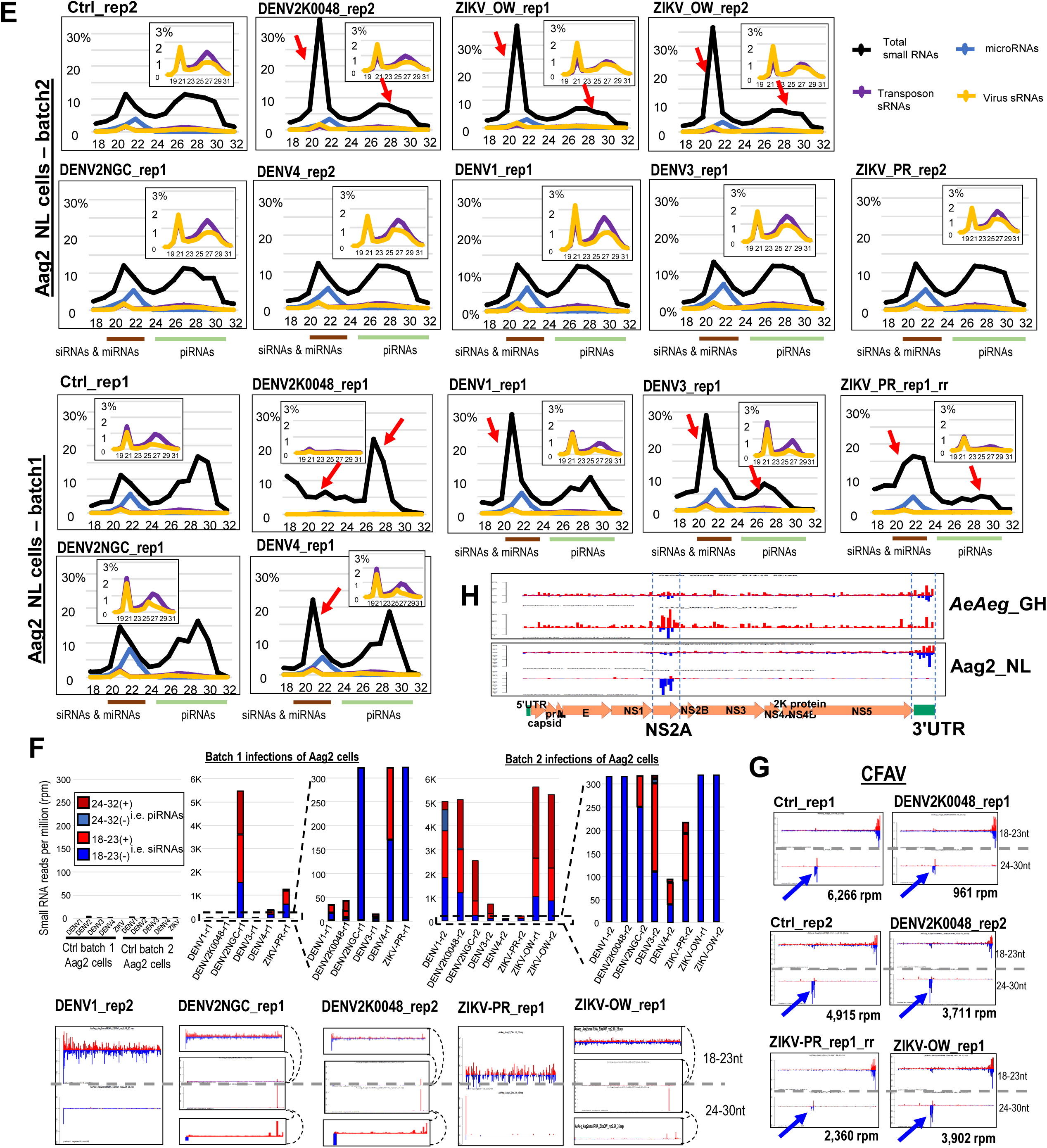
Small RNA profiles from *Aedes aegypti* (*AeAeg*) mosquitoes and cell cultures during viral infections. (A) Small RNA profiles of additional arboviruses in *AeAeg* cell cultures and CHIKV added ectopically to cells from {Morazzani, 2012 #1815} (B) Re-analysis of ZIKV and CFAV small RNAs from *AeAeg* females as sequenced from Saldana et al. Blue arrow notes emergence of new piRNAs from CFAV after active replication of ZIKV small RNAs. To the right of the virus profiles are the small RNA length distribution plots of these libraries. (C) Quantitative RT-PCR assessment of arbovirus replication during two independent batches of infection of Aag2-NL cells. (D) Small RNA functional annotation proportions from the re-analysis of the {Saldana, 2017 #1785} datasets and the two batches of Aag2-NL cell infections with various DENV and ZIKV serotypes. (E) Small RNA length distributions as a proportion of the small RNA library. Inset graph zooms in on the modest proportions of viral and transposons small RNAs. Red arrows point to the significant change from the normal proportion of small RNAs in control cells. (F) Control Aag2-NL cells do not exhibit any small RNAs mapping to DENV and ZIKV, however varying amounts of siRNAs and piRNAs specific to the individual serotypes of DENV and strains of ZIKV are detected in Aag2-NL cells 7 days after infection. Plots below show small RNA mapping patterns to DENV and ZIKV divided by siRNAs versus piRNAs, and insets above and below main plots are zoomed-in narrower Y-scale views of those viruses because in the main view a single plus-strand peak of a very abundant small RNA distorts the view of the other less abundant and distributed small RNAs. (G) Counts and small RNA profiles from CFAV in control and infected Aag2-NL cells. Blue arrows point to pre-existing group of negative strand piRNAs potentially because of multiple pre-existing viruses replicating and generating small RNAs in Aag2-NL cells. (H) The regions generating notable piRNAs and siRNAs from CFAV in mosquitoes and Aag2 cells are the NS2A gene and 3’UTR.

**Figure S7.**
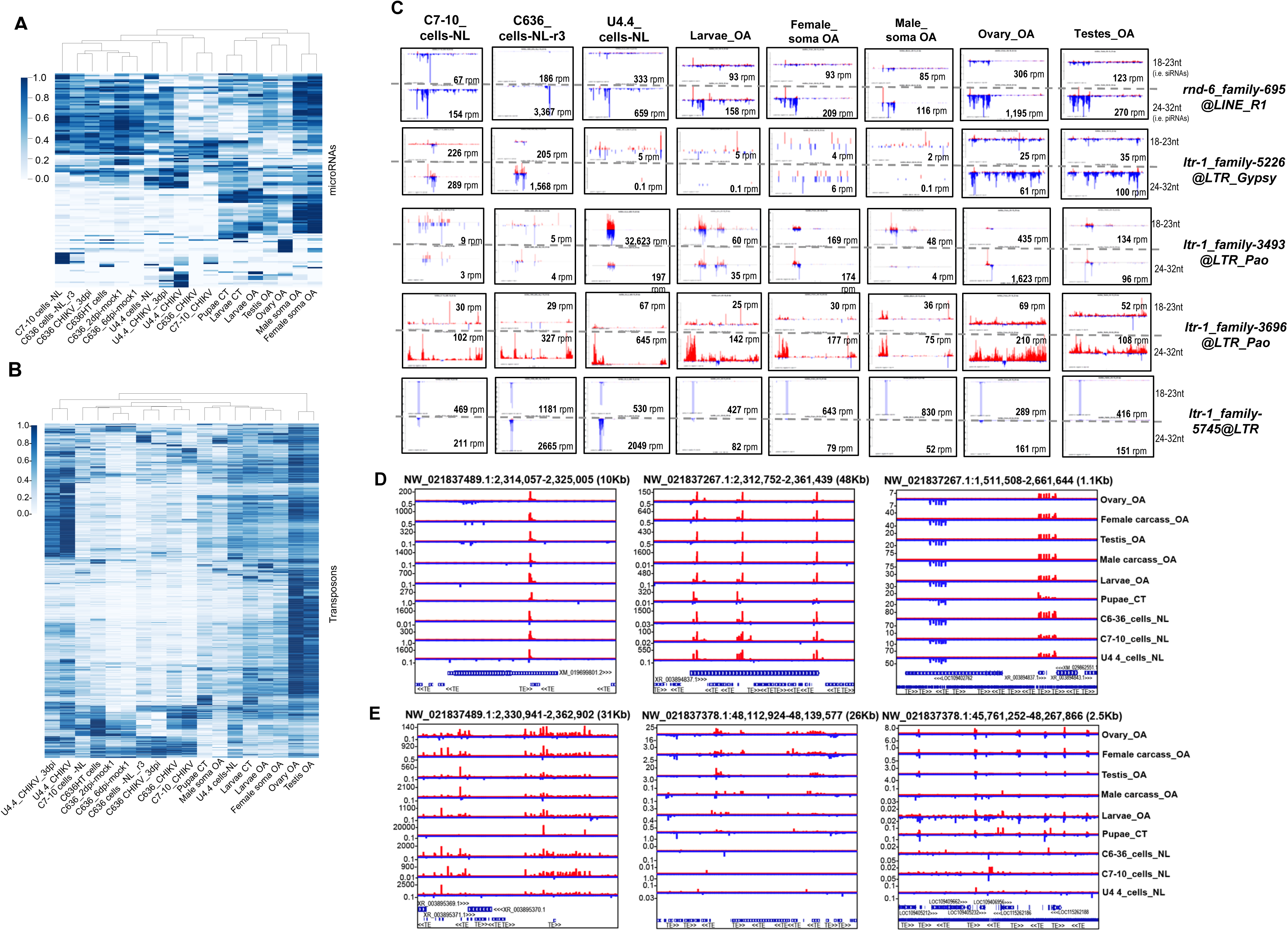
Hierarchical clustering heatmaps of *AeAlbo* microRNA and transposon small RNA profiles, highly expressed transposon small RNA patterns common to cell lines and in the mosquito; and notable *AeAlbo* piRNA clusters. Hierarchical clustering of (A) miRNAs and (B) transposon-mapping small RNA patterns for the cell lines versus the mosquito tissues. (C) Profiles of the *AeAlbo* transposons with most abundant small RNAs both in cell cultures and *AeAlbo* tissues. (D) Genome browser views of notable genic piRNA Cluster Loci (piRCL) in *AeAlbo*. (D) Genome browser views of notable intergenic piRCLs in *AeAlbo*.

**Figure S8.**
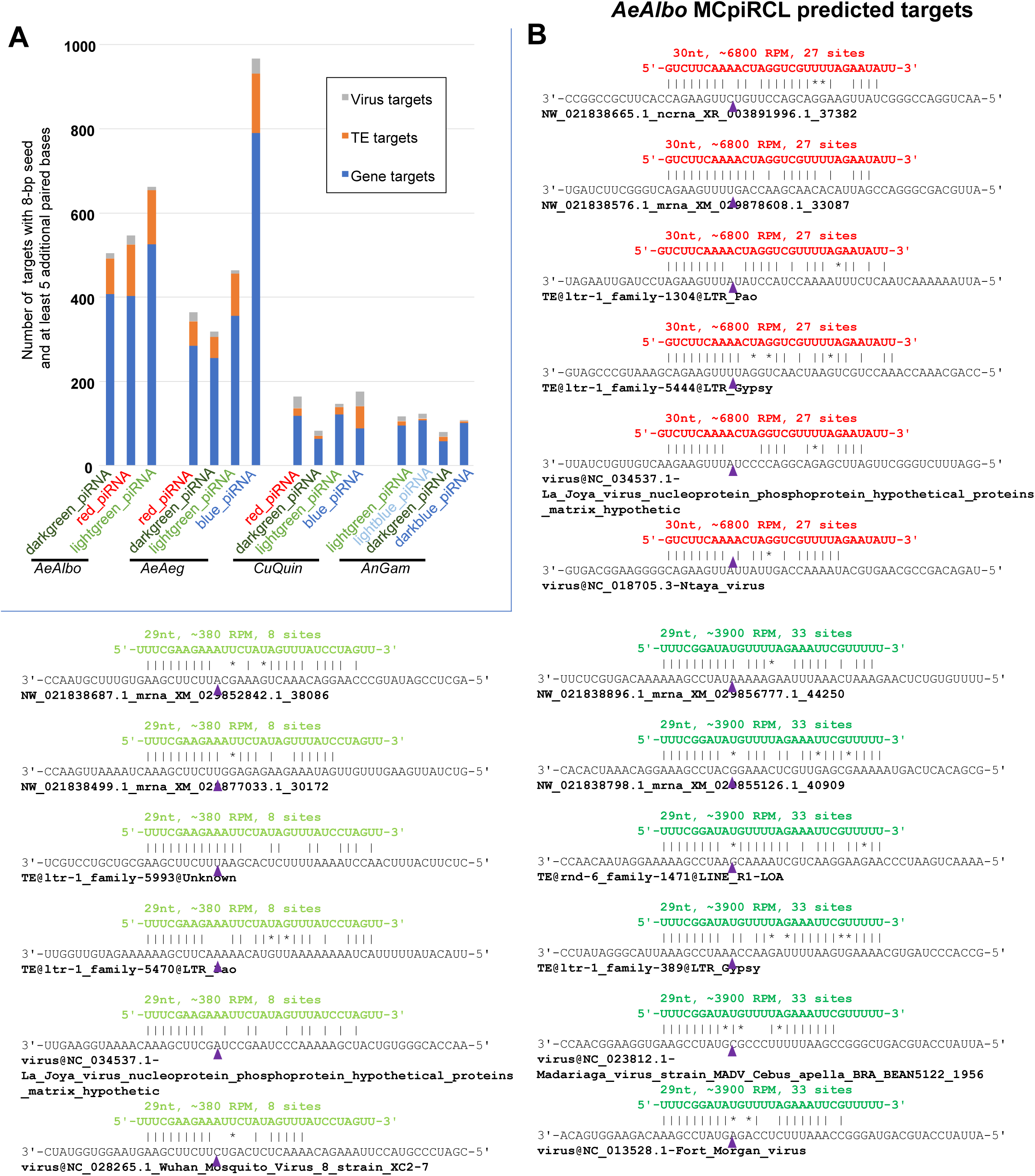
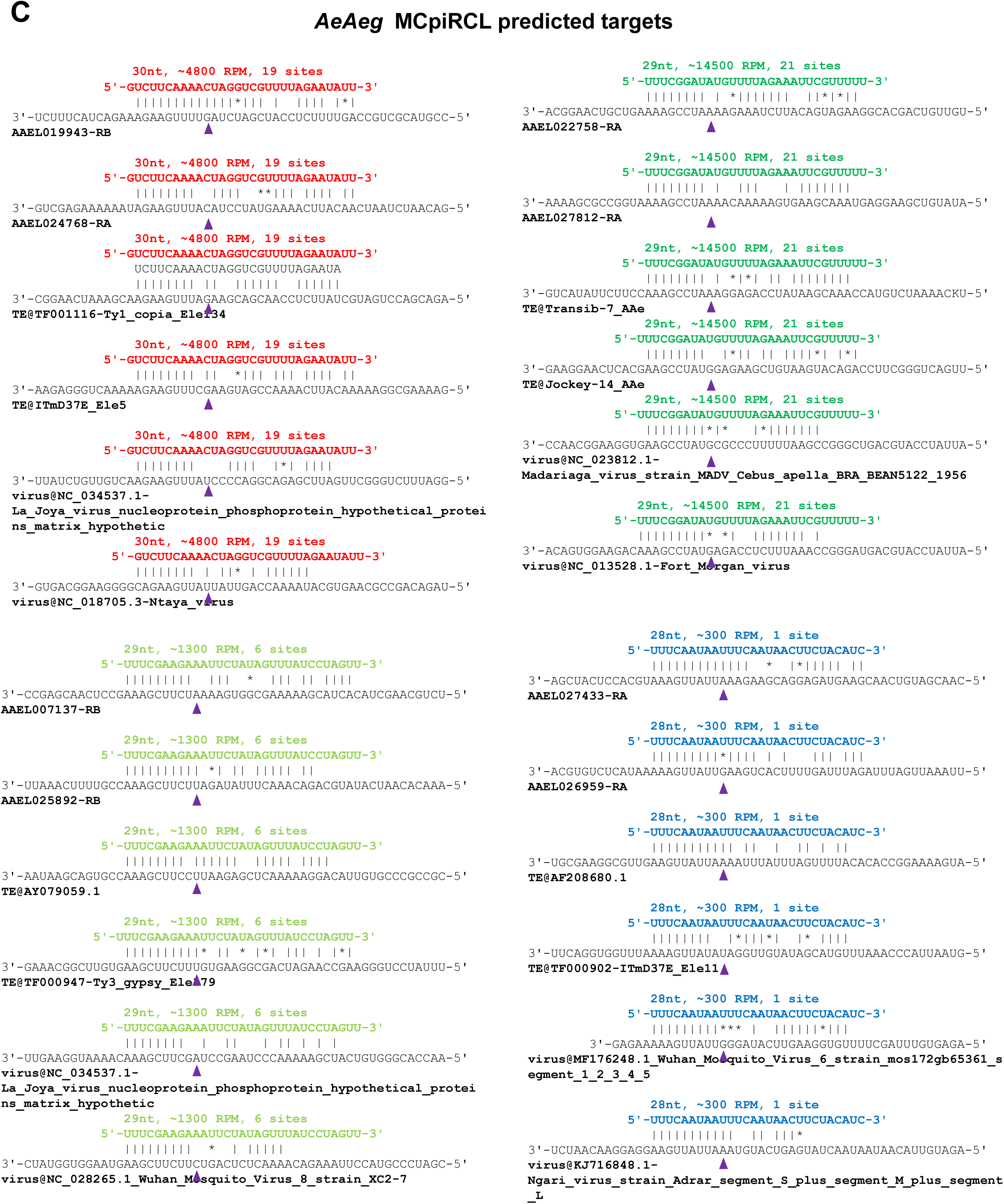
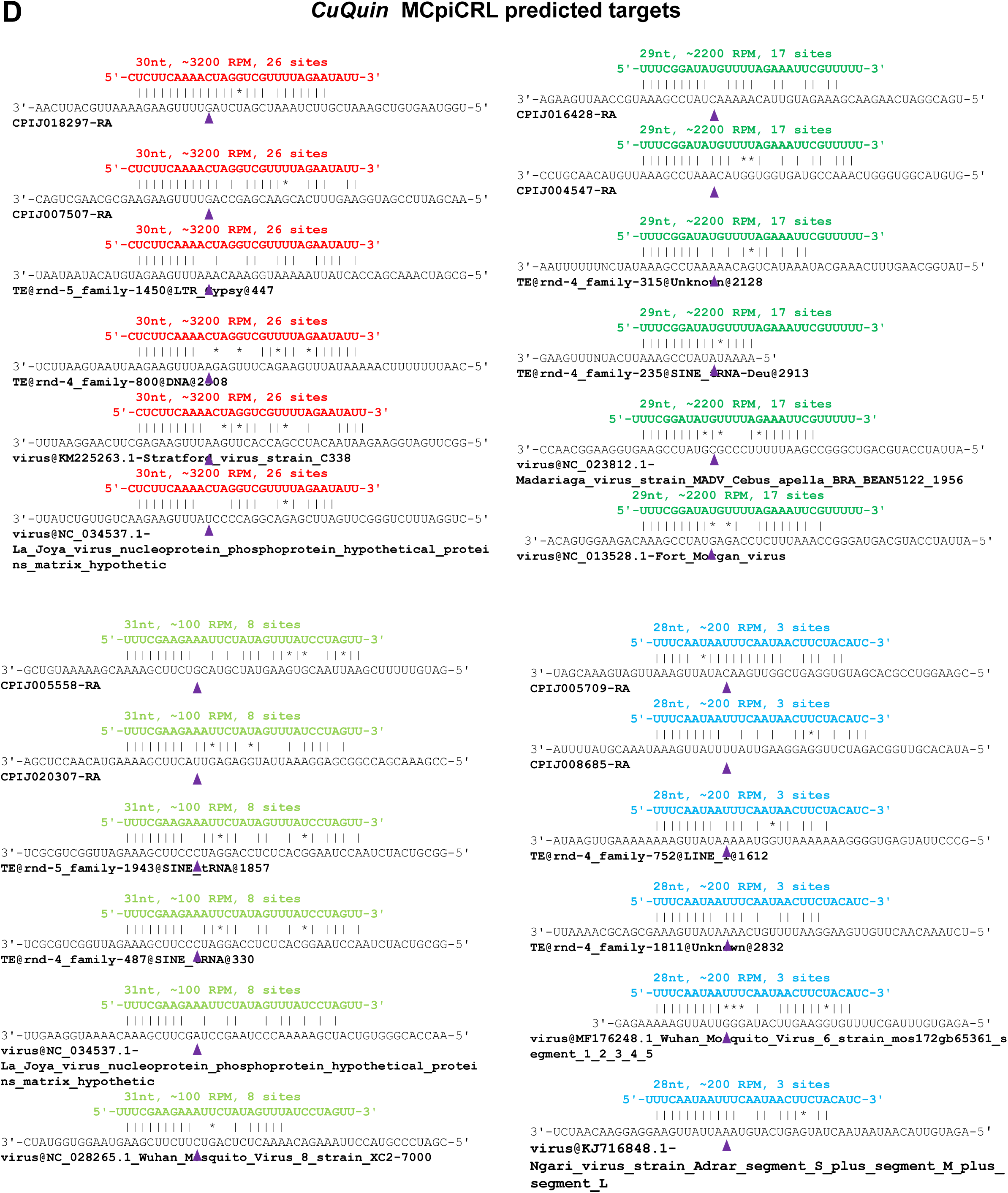
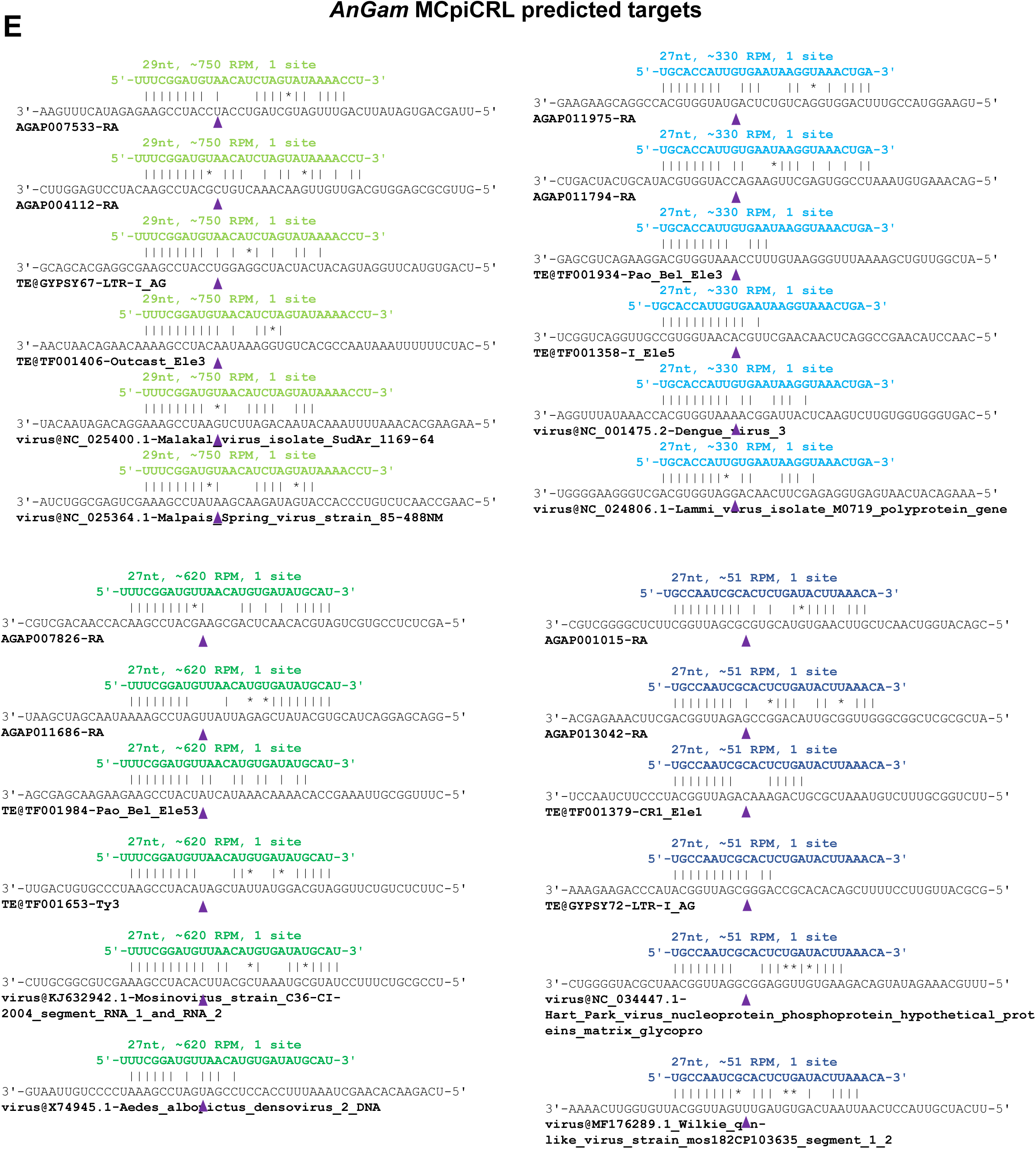
Predictions of gene mRNAs, transposons and virus transcripts potentially targeted by the four piRNAs in the Mosquito Conserved piRNA Cluster Locus (MCpiRCL). (A) Total counts of predicted targets from the four mosquito species. (B) For each of the *AeAlbo* piRNAs from its MCpiRCL, the top two mRNA, transposon and virus target sites is displayed, ranking based on the degree of base-pairing. (C) Top targets for the *AeAeg* piRNAs from its MCpiRCL. (D) Top targets for the *CuQuin* piRNAs from its MCpiRCL. (E) Top targets for the *AnGam* piRNAs from its MCpiRCL.

**Figure S9.**
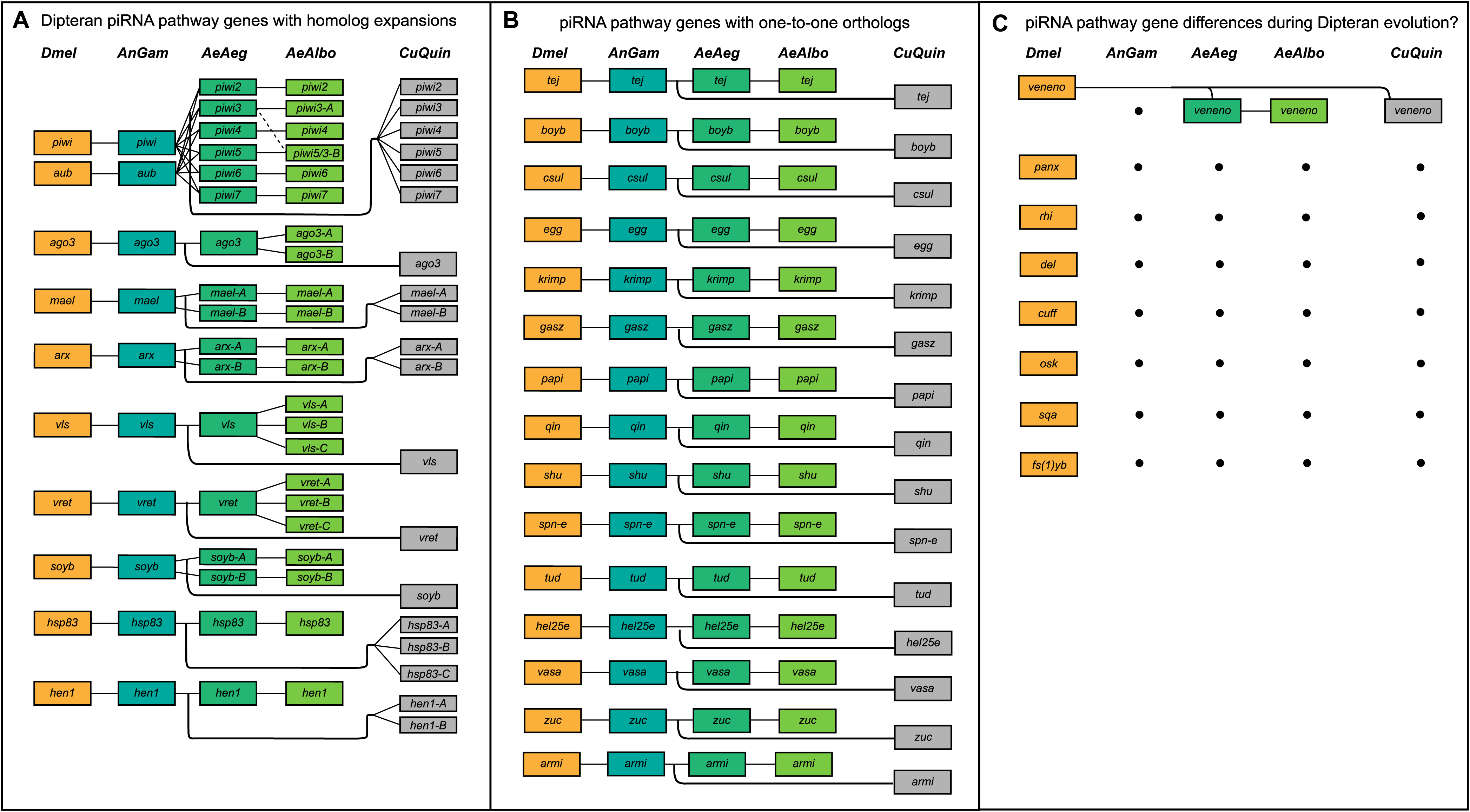
Diagram of the piRNA pathway genes’ conservation between *Drosophila* and mosquitoes. (A) Genes are grouped by the gene sets that have expanded families in culicine mosquitoes. (B) Gene sets that have just one-to-one orthologs. (C) Genes that have been lost in a mosquito lineage.

**Figure S10.**
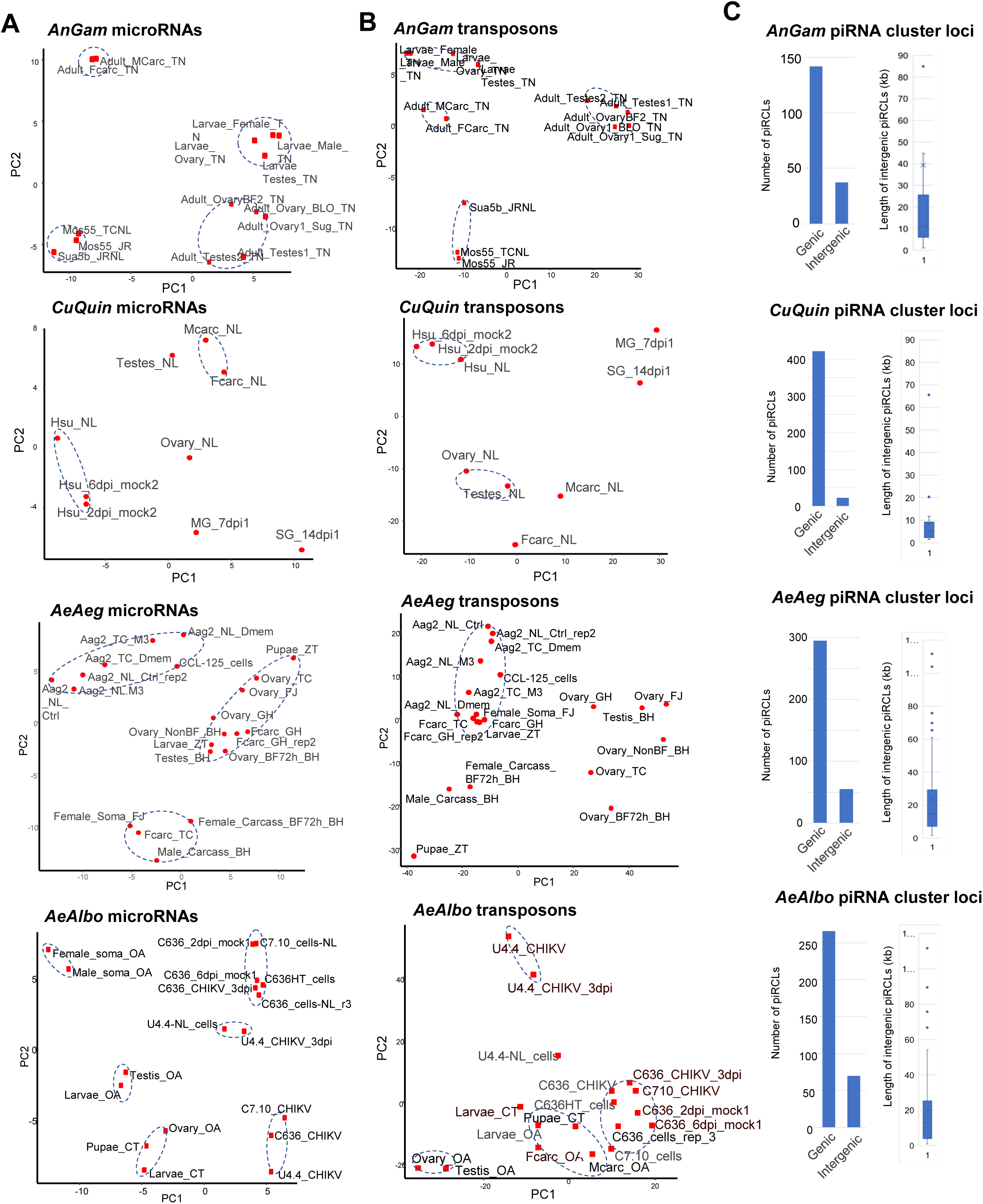
Principal Component Analysis (PCA) of mosquito cell lines and mosquito tissues, and tabulation counts of endogenous piRNA clusters and sizes.

Table S1. Characteristics and culture conditions of the eight mosquito cell lines analyzed in this study. This table only lists the main name of the cell line but the origins of certain cell lines and tissues throughout the study are further delineated with a suffix of initials of the lab providing the mosquito sample.

Table S2. Metatable of *AnGam* samples and cell lines, and additional tabs of top Genic and Intergenic piRNA clusters.

Table S3. Metatable of *Dmel* samples and cell lines re-analyzed for this study for the purpose of investigating piRNA biogenesis patterns.

Table S4. Metatable of CuQuin samples and cell lines, and additional tabs of top Genic and Intergenic piRNA clusters.

Table S5. Metatable of *AeAeg* samples and cell lines, and additional tabs of top Genic and Intergenic piRNA clusters.

Table S6. Metatable of AeAlbo samples and cell lines, and additional tabs of top Genic and Intergenic piRNA clusters.

Table S7. Conservation analysis of homologous genes in the piRNA and miRNA/siRNA pathways in Dipterans.

